# Spatiotemporal Patterns Differentiate Hippocampal Sharp-Wave Ripples from Interictal Epileptiform Discharges in Mice and Humans

**DOI:** 10.1101/2025.02.06.636758

**Authors:** Anna Maslarova, Jiyun N Shin, Andrea Navas-Olive, Mihály Vöröslakos, Hajo Hamer, Arnd Doerfler, Simon Henin, György Buzsáki, Anli Liu

## Abstract

Hippocampal sharp-wave ripples (SPW-Rs) are high-frequency oscillations critical for memory consolidation in mammals. Despite extensive characterization in rodents, their application as biomarkers to track and treat memory dysfunction in humans is limited by coarse spatial sampling, interference from interictal epileptiform discharges (IEDs), and lack of consensus on human SPW-R localization and morphology. We demonstrate that mouse and human hippocampal ripples share spatial, spectral and temporal features, which are clearly distinct from IEDs. In 1024-channel hippocampal recordings from APP/PS1 mice, SPW-Rs were distinguishable from IEDs by their narrow localization to the CA1 pyramidal layer, narrowband frequency peaks, and multiple ripple cycles on the unfiltered local field potential. In epilepsy patients, ripples showed similar narrowband frequency peaks and visible ripple cycles in CA1 and the subiculum but were absent in the dentate gyrus. Conversely, IEDs showed a broad spatial extent and wide-band frequency power. We introduce a semi-automated, human ripple detection toolbox (“ripmap”) selecting optimal detection channels and separating event waveforms by low-dimensional embedding. Our approach improves ripple detection accuracy, providing a firm foundation for future human memory research.

## Introduction

Hippocampal sharp-wave-ripples (SPW-Rs) are essential for the compression, chunking, and delivery of neural codes to the neocortex^1,2^. Decades of rodent research have established the role of SPW-Rs in consolidating and guiding experience. In rodents, place cell sequences acquired during exploratory behavior are replayed in SPW-Rs during offline states, such as consummatory behaviors and NREM sleep^3–5^, when acetylcholine levels are decreased^6^. SPW-Rs trigger widespread neocortical activation potentially inducing long-term synaptic plasticity in hippocampal output regions^2,5,7–12^. Selective prolongation or erasure of SPW-Rs leads to respective memory enhancement or impairment during rodent spatial navigation, supporting their essential role^4,13,14^.

In rodents, hippocampal electrodes with high spatial precision confidently detect SPW-Rs and their corollary spike bursts. SPW-Rs appear in the local field potential (LFP) as a negative sharp-wave in the CA1 stratum radiatum and brief high-frequency oscillations (100-200 Hz) in the CA1 pyramidal layer. CA3 and CA2 population bursts drive coordinated oscillatory excitation of CA1 interneurons and pyramidal cells, underlying the fast ripple oscillation^15–17^. In the CA dendritic layers and the dentate gyrus (DG), the LFP is dominated by a negative sharp wave. The strong excitatory output from CA1 during SPW-Rs can induce lower frequency but similar duration ripples in the target subicular complex, retrosplenial cortex and deep entorhinal cortex layers^12,18–20^.

Recent human studies report hippocampal ripples during NREM sleep and awake tasks^21–27^ occurring at lower frequencies than rodent SPW-Rs^24,25,28–30^. Unlike rodent SPW-Rs, which reflect synchronous firing in the CA3/CA1 circuit, human ripples are broadly defined as “ripple band” or high-frequency oscillations (HFOs) in the hippocampus and neocortex and their sharp-wave component is rarely reported^30–32^. Most approaches use a simple bandpass filter (between 80-120 Hz^25,28,31,32^ or up to 200 Hz^24,26,29,33^) to detect brief (30-100 ms) events above a certain threshold^24,31^. Ripples have been detected during NREM sleep, phase-coupled with spindles and slow oscillations^23,28^, similar to rodents^34^. Yet, ripples during awake, effortful memory processes have also been reported, with increased ripple rates during successful learning and retrieval^21,24,25,33^. Hippocampal ripples during autobiographical recall have been demonstrated to trigger widespread increases in high gamma activity in regions associated with default mode network (DMN)^35^.

These studies suggest a translational link between human ripples and rodent SPW-R activity. However, given the poor spatial resolution and sparse sampling of depth electrodes in the human hippocampus, contamination with IEDs in epileptic brain tissue, muscle artifacts, and lack of arousal state measurements, separation of SPW-Rs from other physiological (e.g., high gamma) and pathological HFOs is fraught (for a review see^29^). Clinical macroelectrodes typically sample from tens of thousands of neurons^36^, and rarely target the CA1 subfield^24,35^. Confirmation of electrode placement to the CA1pyramidal layer, standard in rodents, is difficult to achieve and not routinely attempted in human studies. The large size of the macroelectrode (∼1mm diameter) would likely span several subfields and layers during surgical implantation. Finally, detection parameters across human studies vary widely, resulting in a wide distribution of ripple incidence across sleep, from 0.35 to 30 ripples per minute^22,29^. The practical constraints of clinical macroelectrode recordings and variable detection features across human literature raise uncertainty in the validity of ripple reports in human cognitive studies and their cross-species translation.

We address these challenges by comparing high-resolution hippocampal recordings from all hippocampal subfields in an Alzheimer’s disease (AD) mouse model to macro- and microwire recordings in surgical epilepsy patients. The AD model, characterized by hippocampal IEDs with rare seizures, was chosen to mimic the human interictal brain state^37,38^. Using wide-coverage 1024-channel probes (SiNAPS)^39,40^, we captured rodent SPW-R activity across all hippocampal subfields and layers simultaneously enabling precise characterization of spectrotemporal features. Simultaneous IED sampling allowed us to define distinct features of physiologic and pathologic events regardless of recording location. With these guiding features, we identified putative ripples in macro- and microwire recordings in 14 surgical epilepsy patients. Manual curation discarded 90% of automatically detected ripples as false positives. From this curated dataset of high-confidence ripples, we identified parameters to optimize human ripple detection on macrocontacts and microwires. Finally, we developed a Uniform Manifold Approximation and Projection (UMAP)-based semi-automatic separation approach and toolbox (“ripmap”) to discard IEDs and false-positive events, reducing the need for manual curation by up to 45%.

## Results

### Challenges of human SPW-R detection compared to rodents and important definitions

In rodents, high-density probes can simultaneously record across hippocampal subfields and cellular layers at the micrometer scale (Fig. 1a), and their positions can be validated by histology (Fig.1b). Moreover, characteristic LFP features on laminar recordings from the hippocampus enable real-time identification of recording regions. The distinct laminar morphology of SPW-Rs facilitates differentiation from other high-frequency events, such as IEDs (Fig. 1c). In contrast, human probes sample signals at the millimeter scale and lack the density of rodent probes (Fig.1d). Recording positions are estimated by visualizing contacts on a postoperative computer tomography (CT) - scan, aligned to the patient’s magnetic resonance imaging (MRI) data, and electrode coordinates transferred to the MRI (Fig. 1e). Microwires typically used for sampling of hippocampal single neurons, spread arbitrarily across hippocampal layers and subfields, lacking laminar specificity (Fig. 1f). This precludes the reliable use of LFP features to identify individual channel positions. Signal analysis is further complicated by high-power IEDs resembling physiological oscillations in some regions (Fig.1f) and short-lived “high-gamma” oscillations overlapping with ripple frequencies.

**Fig. 1.**
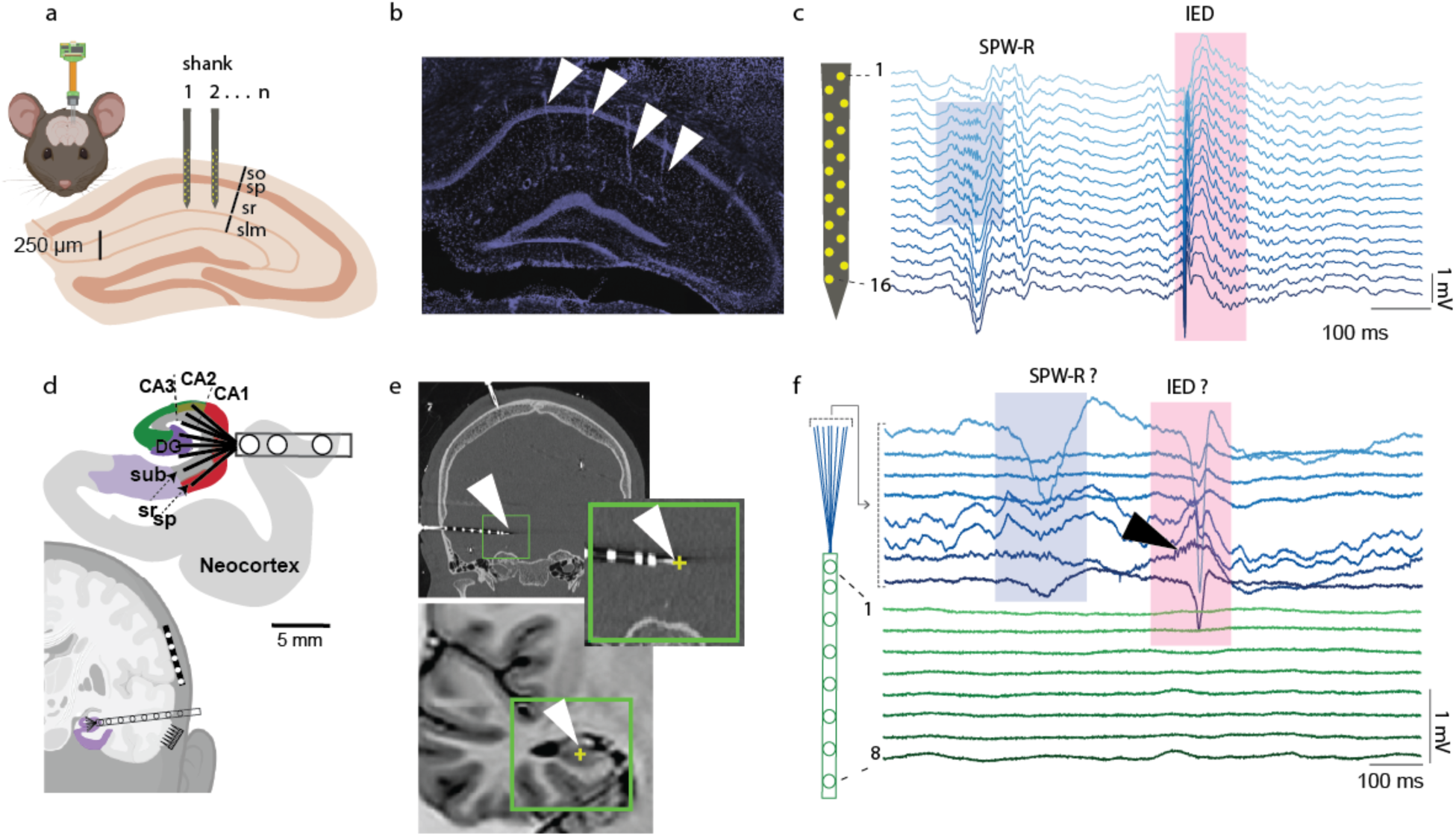
Comparison to standard recordings in rodent hippocampus reveals shortcomings in human recording methods: **a Top left**: Multi-shank, high-density probe implanted in the mouse hippocampus. **Bottom:** Close-up on the recording probe with contacts (in yellow) covering several CA1 layers: so - stratum oriens, sp – stratum pyramidale, sr – stratum radiatum, slm – stratum lacunosum-moleculare. Two shanks are displayed. **b** Validation of shank position in rodents: DAPI stained coronal section of the dorsal hippocampus, as outlined in (a), shows visible traces of 4 probe shanks (white arrows) in the mouse brain. **c** Example mouse local field potential (LFP) recording from one shank of a 64-channel probe (4×16, NeuroNexus) with the probe’s tip in CA1. A sharp wave-ripple (SPW-R; blue box) is detected in the CA1 pyramidal layer but not above the hippocampus. One interictal epileptiform discharge (IED; red box) shows a distinct laminar distribution and is detected on all channels. **d Bottom:** Conventional methods for LFP and single-unit sampling in surgical patients: cortical grid electrode and a cortical multielectrode array over the right neocortical structures, a depth macro/microwire probe in the left temporal lobe **Top:** Close-up on the tip contacts of a Behnke-Fried macro/microwire depth electrode implanted in the temporal lobe. The flexible microwires spread arbitrarily. In this example, the microwires are shortened at 5 mm and scaled to a hippocampus image from the Allen Brain Atlas to demonstrate possible coverage of distant hippocampal subfields. Three macrocontacts (1mm) are sampling adjacent neocortical structures. **e** Contact localization in humans. Left image: postoperative coronal CT section of a patient implanted with depth electrodes. The microwire is visible at the tip of the electrode (white arrow and inset). Right image: preoperative T1 MRI coronal section corresponding to the CT section from the same patient. The MRI was co-registered with the postoperative CT, and the coordinates of the microwire tip were transferred from the CT to the MRI scan (yellow cross) to estimate the anatomical location of the wire. **f** Example LFP recordings from a human depth electrode with 8 microwires (blue), and 8 macro-contacts (green). The order of the microwires is arbitrary. One putative SPW-R event (blue box) precedes a more ambiguous, IED-like, event (pink box), which resembles a SPW-R on channel 7 – black arrow.

To compare events in rodents and humans, we adopt the following terminology: sharp-wave ripples, or SPW-Rs, for events detected on CA1 pyramidal layer channels and high-frequency oscillations, or HFOs, for events on non-CA1 channels in rodents. Because human recordings are coarser and rarely detect the sharp wave component, we refer to putative sharp-wave ripples as “**ripples**.” Other high-frequency oscillatory and aperiodic detections, including gamma or pathological high-frequency oscillations, we refer to as **false positive events**.

### Rodent SPW-Rs are localized to CA1 pyramidal layer

Automatic SPW-R detection by conventional band-pass filtering for the high-frequency oscillation can lead to false positives if the detection is performed in regions remote from CA1. To evaluate the spatial, temporal, and morphological features of SPW-Rs across the hippocampus, we utilized high-density recordings spanning the entire dorsal hippocampus in APP/PS1 mice using 1024 channel 8-shank probes (SiNAPS, NeuroNexus and Corticale SRL, Fig. 2a). This allowed us to view SPW-Rs across all hippocampal subfields simultaneously (Fig. 2b). SPW-Rs were confined to the CA2/CA1 axis, consistent with earlier findings using lower-density probes^41,42^. During CA1 SPW-Rs, CA3 exhibited population spike bursts and sharp waves with less prominent and variable ripples, while the dentate gyrus displayed irregular negative waves on event-averaged traces (Fig. 2c). Current source density (CSD) analysis confirmed that SPW-Rs resulted in current sinks in the CA1 stratum radiatum and sources in the pyramidal layer. Wavelet spectrograms showed a narrow ∼160 Hz peak in CA1 pyramidal and dendritic layers, absent in CA3 and dentate gyrus, where low frequency gamma events (frequency distribution peaks ∼ 20 Hz) predominated, possibly reflecting filtered versions of SPW-Rs (Fig. 2d).

**Fig. 2.**
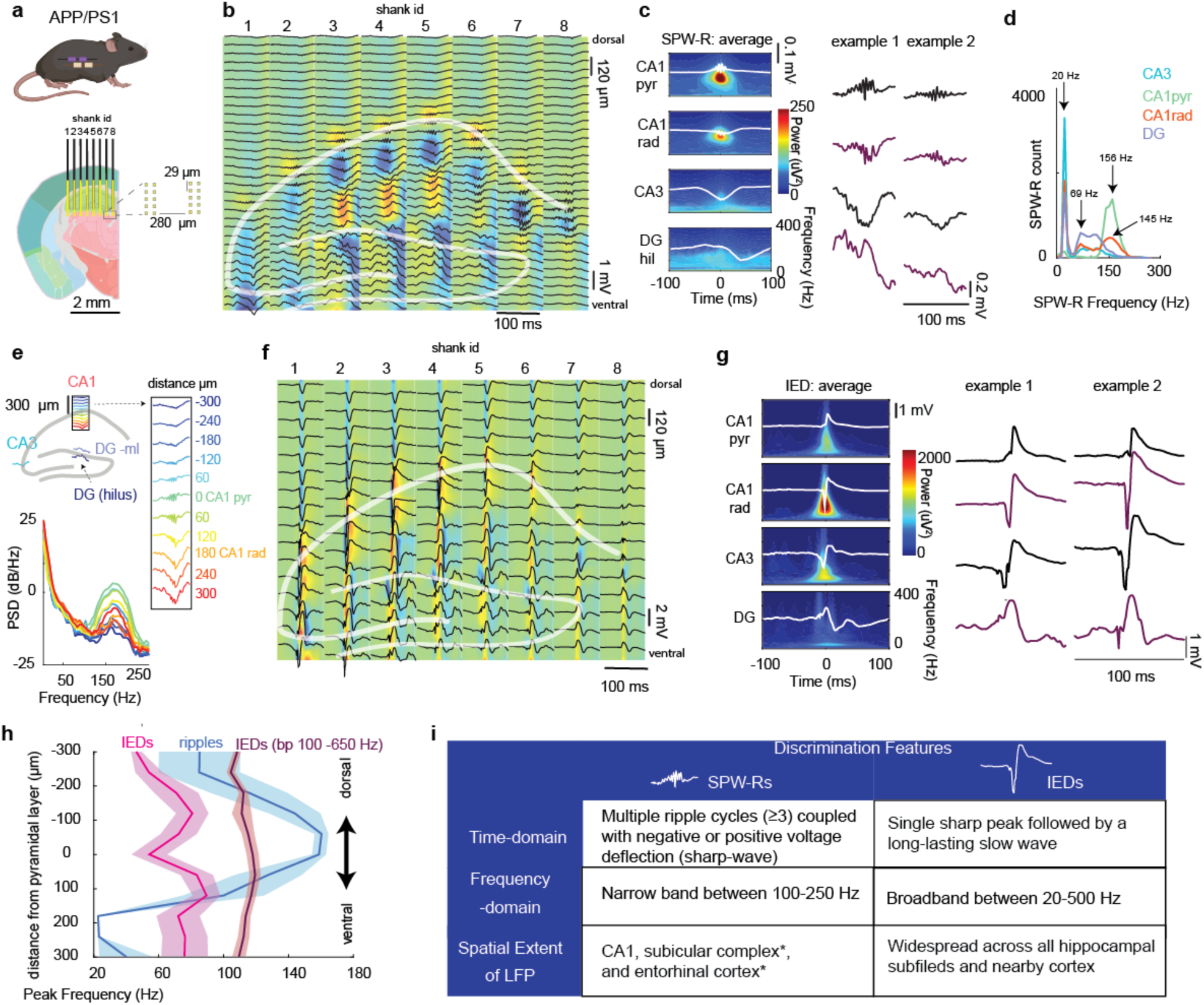
Mouse hippocampal SPW-Rs and IEDs display distinct spatial, spectral and morphologic features. **a** Recording configuration of an 8-shank, 1024 channel probe (SiNAPS, NeuroNexus and Corticale Srl), positioned in the dorsal hippocampus of an APP/PS1 mouse. The dimensions of the probe enable sampling of all hippocampal subfields and neighboring deep cortical layers. Inset: schematic of the pixel design of the probe – two channels per row and shank. Inter-shank distance and inter-contact distances are indicated. **b** Example recording from a head-fixed APP/PS1 mouse displays a SPW-R detected in the CA1 pyramidal layer. LFP traces of every 4th channel are shown and plotted on top of an average current source density map from all SPW-Rs in one recording session (n= 803 SPW-Rs). The CA layers and granule cell layer are outlined in white. **c** Average LFP traces (white) and wavelet spectrograms from the CA1 stratum pyramidale (pyr), CA1 stratum radiatum (rad), CA3 pyramidal layer and hilus of the dentate gyrus (DGhil). Ripples are observed only in CA1. Right: two example SPW-R events. **d** Histograms of the peak frequencies of CA1 SPW-Rs on the same channels as in (b). **e** Above: Schematic of the locations of channels used for analysis in plots c, d, g and h: CA1 - pyramidal channel, selected by the highest high-frequency power (100-500 Hz). The black rectangle shows channels within 300 µm from the CA1 pyramidal channel, defined as distance 0 in the power-spectral density plot (PSD) plot below and in (h). Below: Peri-SPW-R PSD at different depths. **f** Example IED event detected on a CA1 pyramidal channel, LFP traces of every 8th recording channel are plotted on top of the current source density map, calculated from IEDs in one recording session (n=7 IEDs). **g** Average LFP traces (white) and wavelet spectrograms of IEDs from one recording (n=13 IEDs). IEDs were detected in CA1 rad. Corresponding traces are shown from CA1 pyr, CA3 and DG hilus. Right: two example IEDs. **h** Peak frequency of SPW-Rs (ripples, blue) and IEDs (pink), as a function of channel distance from the CA1 pyramidal layer (distance 0, x-axis). Negative values indicate channels above the pyramidal layer (ventral, towards cortex), as shown in (e). The purple curve represents peak frequency of IEDs on band-pass filtered data (bp 100-600 Hz). **i** Summary of distinctive SPW-R and IED features defined from the mouse hippocampus. * indicates regions not tested in this study but reported in other rodent studies.

To quantify the spatial extent of SPW-Rs, we calculated the peak frequencies of the signal during SPW-Rs, as a function of distance from CA1 pyramidal layer (Fig. 2e). Peak frequencies were highest in the CA1 pyramidal layer (160±4 Hz, mean/SEM) and the peak power in the ripple band rapidly decreased with distance. Peak frequencies also decreased to 85±25 Hz in stratum oriens, and 40±17 Hz in stratum lacunosum-moleculare (n=9, Fig. 2h).

To determine whether HFOs detected in other hippocampal subfields reflect CA1-SPW-R activity, we used the same detection parameters to identify HFOs in CA3, dentate gyrus molecular layer and hilus independently, then measured their temporal coupling to CA1-SPW-Rs (Supplementary Fig. 1). These HFOs were weakly temporally coupled to CA1 SPW-Rs (±200 ms window, CA3: 17±5 %, n=7, DG hilus: 8±1 %, DG molecular layer: −8±2 %, n=9, Supplementary Fig. 1g). Mean peak frequencies of HFOs in CA3 (103±0.5 Hz) and dentate gyrus (106±0.3 Hz) were higher than SPW-Rs detected in CA1 (CA3: 57±0.7 Hz; DG: 82±0.5 Hz, Supplementary Fig. 1a-c, d, h, p < 0.0001, Wilcoxon rank sum test). We next compared the frequency, duration and amplitude of HFOs coupled with CA1 SPW-Rs vs uncoupled HFOs (Supplementary Fig. 1d-f). Coupled CA3 HFOs showed slightly higher peak frequencies and longer durations. In the DG, coupled HFOs showed lower amplitudes but longer durations.

Finally, we examined whether specific features of CA1 SPW-Rs predict coupling with HFOs in other regions. Coupled SPW-Rs had slightly higher peak frequencies, larger amplitudes and longer duration, compared to uncoupled events (Supplementary Fig. 2). In summary, >90% of HFO events detected in hippocampal subfields outside of CA1 were not temporally coupled to CA1 SPW-Rs and possessed differing spectrotemporal features. These findings suggest that SPW-Rs are tightly localized to the CA1 subfield, and that HFO events outside CA1 are likely false positives.

### Distinct features of SPW-Rs and IEDs

IEDs are pathological hypersynchronous population bursts observed between seizures^43–45^. Electrographically, IEDs are transient events clearly distinguished from the background, lasting 20-70 ms (“spike”) or 70-200 ms (“sharp wave”), followed by a negative slow wave. IEDs may be phase-coupled with pathological HFOs within the mesial temporal lobe seizure-onset zone^46^. Like SPW-Rs, hippocampal IEDs are more frequent during NREM sleep ^47–49^ and appear to propagate along hippocampal efferent pathways ^37,50,51^, possibly through the excessive synchronization of sharp wave events ^44,52^.

To differentiate SPW-Rs from IEDs, we detected IEDs in APP/PS1 mice and characterized their waveforms and frequencies across hippocampal subfields. IEDs showed varying appearances, but the majority were characterized by brief, large-amplitude LFP discharges exceeding SPW-R magnitudes by over five-fold. IEDs were observed in all hippocampal regions (Fig. 2f). In CA1 and CA3 dendritic layers, IEDs typically presented as a negative spike, followed by a slow wave whereas in cell layers, the negative spike was less prominent (Fig. 2f,g). A sharp negative population spike is unique to IEDs^44^, as it was never seen with CA1 SPW-Rs. CSD analysis showed an initial current sink in the DG^53^, with propagation along the CA axis (Fig. 2f). Unlike SPW-Rs, IEDs lacked ripple-like oscillations but exhibited a broad-band (20-400 Hz) power increase, approximately 10-fold higher than SPW-Rs, possibly reflecting the large-amplitude sharp spike (Fig. 2g). IED peak frequencies were lower than SPW-Rs (54±11Hz in the pyramidal layer) and remained similar with distance from CA1 pyramidal layer (300 µm above: 46 ±3 Hz, 300 µm below: 76 ±14 Hz). IED peak frequencies calculated from filtered signals, typically used for peak frequency extraction, were somewhat higher (pyramidal layer: 118±4 Hz, 300 µm above: 108±2 Hz, 300 µm below: 109±4 Hz, Fig. 2h). These peaks likely reflect edge filtering artifacts, as no ripple oscillations were visible in the raw LFP during IEDs. Thus, HFOs visible on filtered LFP and multiple subfields with broadband peak frequencies likely represent IEDs.

In summary mouse SPW-Rs and IEDs showed distinct LFP morphology (SPW-Rs – multiple ripple cycles; IEDs - a sharp spike and wave), spectral features (narrow-band vs broad-band high-power increase) and spatial distribution (CA1 pyramidal layer vs all hippocampal regions and layers, Fig. 2i).

### Rodent findings guide human SPW-R detection and demonstrate consistency with expert review

Key SPW-R and IED features identified in mice (Fig. 2i) were applied to classify ambiguous LFP waveforms detected in humans as ripples, IEDs, and other false positive events. We analyzed hippocampal recordings from Behnke-Fried macro-microwire hybrid electrodes (Fig. 3a and Fig. 1d,f) in 14 patients during surgical evaluation (6 temporal lobe epilepsy; 7 frontal lobe epilepsy, 1 with occipital lobe epilepsy, Table 1). Ripple detection was conducted across hippocampal macrocontacts and microwires using a published automated method by excluding pre-detected IEDs ±1s to avoid high-power artifacts (Supplementary Fig. 5) and then thresholding the band-pass filtered signal (80 to 250 Hz) during sleep. We then visually inspected all detected events to identify electrodes with putative ripple events. Initial automated detection yielded a high ratio of events that did not meet our predefined criteria for ripples determined from rodent recordings (Fig. 2i), or resembled IEDs. We classified these events as either false positives or IEDs (Fig. 3b, gray boxes).

**Fig. 3.**
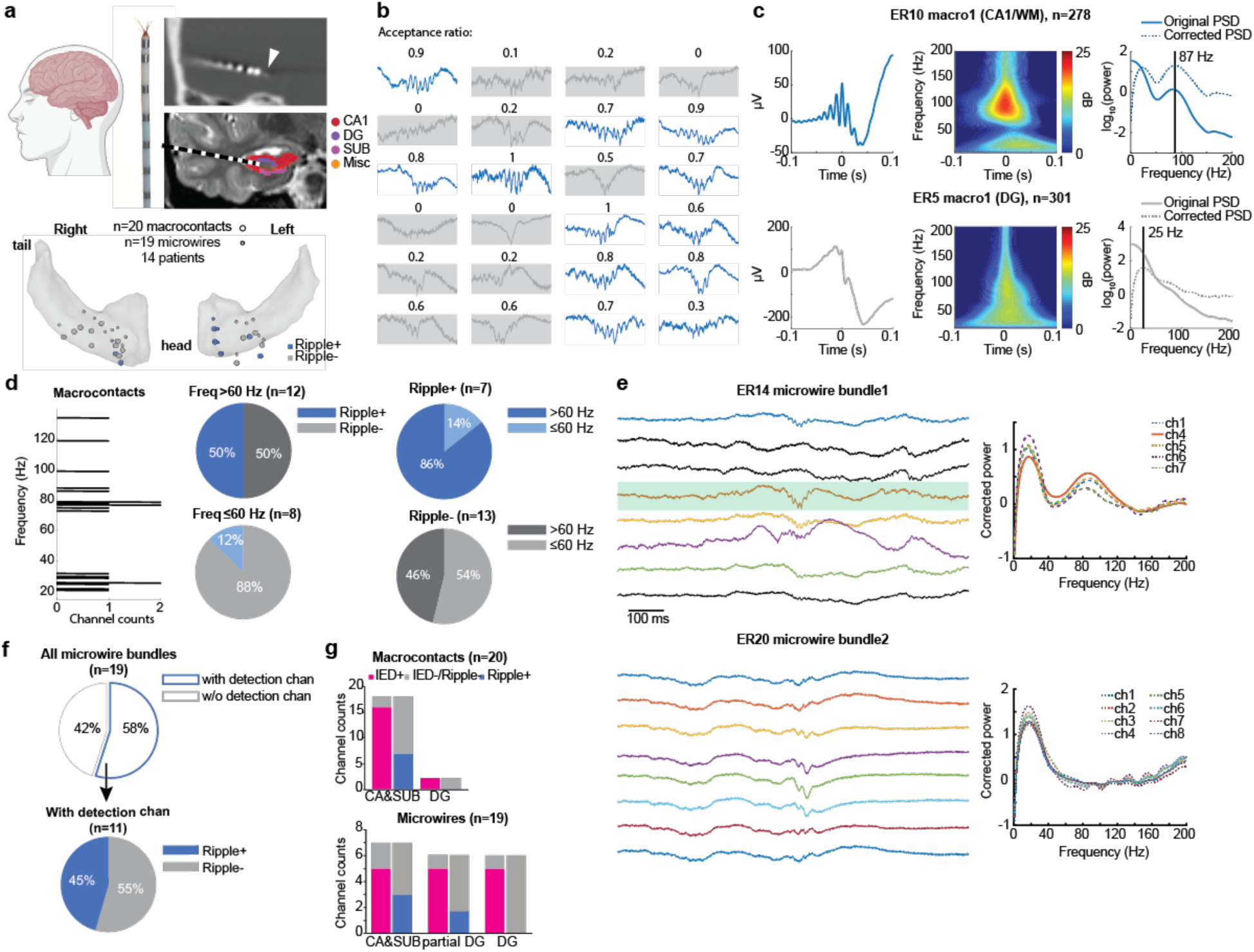
Human ripples are detected in CA/subiculum but not in DG. **a** Top, CT image showing electrode contacts and T1 image overlaid with hippocampal subfields. Arrow: microwires. Left, Behnke-Fried hybrid electrodes. Bottom, Anatomical distribution of electrodes used in this study. Large circles: macro-contacts; small circles: microwire bundles. Blue: Ripple-positive channels. **b** Example manual curation and survey result of a patient data. Grey boxes highlight events that were rejected by visual inspection using the criteria in Fig. 2i. Numbers on top show the acceptance ratio of 10 independent evaluators (0 - no evaluator rated this event as a ripple, 1 - all evaluators rated this event as ripple). **c** Average peri-ripple LFP, wavelet spectrogram and power spectral density (PSD) of example CA1 (top) and DG (bottom) electrodes. Spectral peaks were extracted from 1/f-corrected PSD (dotted lines). **d** Peak frequency distribution of all macro-contacts (left). Percentage of Ripple+ and Ripple-channels (middle). Percentage of channels exhibiting >60 Hz or <= 60 Hz peaks (right). Ripple+ channels predominantly exhibited >60 Hz peaks, while those without ripples showed peaks <60 Hz. **e** Microwire channel selection for ripple detection. Detection channels were selected based on the spectral peak in the ripple band in event-PSDs (upper). When multiple channels showed ripple band peak, the one with the strongest power was selected (green shade). If no channel showed a peak in the ripple band, the entire bundle was considered ripple negative (lower). **f** Percentage of microwire bundles with and without any detection channel (upper). Among 55% or 11 channels with at least one detection channel, 45% or five channels showed ripples (lower). **g** Subfield-specific occurrence of IEDs and ripples. IEDs were observed across all hippocampal subfields, whereas ripples were predominantly found outside of DG.

**Table 1.**
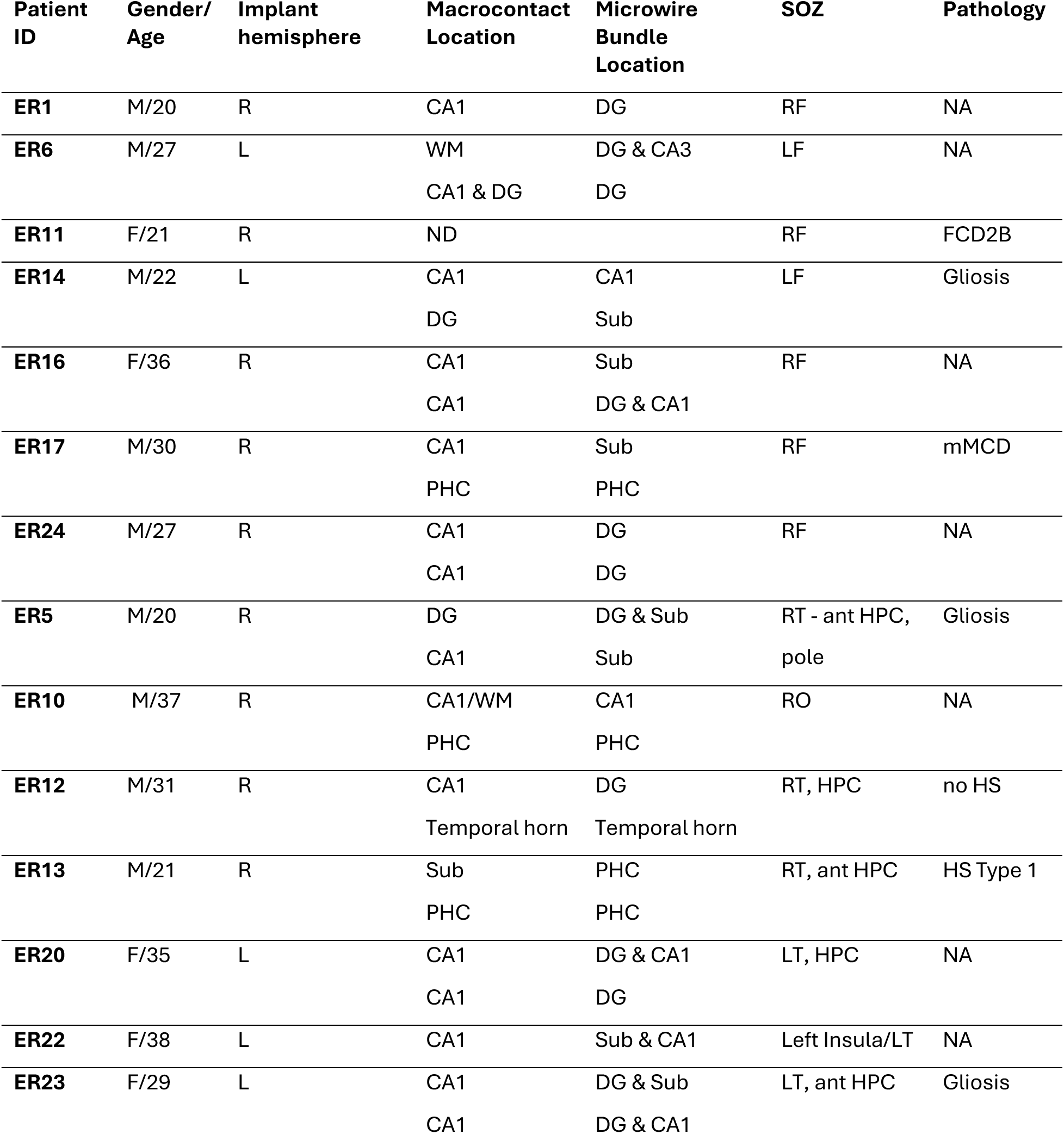
Patient demographics and electrodes. Patients with frontal epilepsy are listed first. When there were two temporal electrodes in a patient, the upper cells indicate the more anterior electrodes. Grey: excluded channels. Legend: F– female, M – male, R – right, L-left, HS – hippocampal sclerosis, NA – not applicable (no resection performed), ND – no data, hippocampal segmentation failed – excluded from analysis, WM – white matter, F – frontal, T – temporal, O – occipital, HPC – hippocampus, FCD – focal cortical dysplasia, mMCD – mild malformation of cortical development

| Patient ID | Gender/ Age | Implant hemisphere | Macrocontact Location | Microwire Bundle Location | SOZ | Pathology |
| --- | --- | --- | --- | --- | --- | --- |
| ER1 | M/20 | R | CA1 | DG | RF | NA |
| ER6 | M/27 | L | WM<br>CA1 & DG | DG & CA3<br>DG | LF | NA |
| ER11 | F/21 | R | ND |  | RF | FCD2B |
| ER14 | M/22 | L | CA1<br>DG | CA1<br>Sub | LF | Gliosis |
| ER16 | F/36 | R | CA1<br>CA1 | Sub<br>DG & CA1 | RF | NA |
| ER17 | M/30 | R | CA1<br>PHC | Sub<br>PHC | RF | mMCD |
| ER24 | M/27 | R | CA1<br>CA1 | DG<br>DG | RF | NA |
| ER5 | M/20 | R | DG<br>CA1 | DG & Sub<br>Sub | RT - ant HPC,<br>pole | Gliosis |
| ER10 | M/37 | R | CA1/WM<br>PHC | CA1<br>PHC | RO | NA |
| ER12 | M/31 | R | CA1<br>Temporal horn | DG<br>Temporal horn | RT, HPC | no HS |
| ER13 | M/21 | R | Sub<br>PHC | PHC<br>PHC | RT, ant HPC | HS Type 1 |
| ER20 | F/35 | L | CA1<br>CA1 | DG & CA1<br>DG | LT, HPC | NA |
| ER22 | F/38 | L | CA1 | Sub & CA1 | Left Insula/LT | NA |
| ER23 | F/29 | L | CA1<br>CA1 | DG & Sub<br>DG & CA1 | LT, ant HPC | Gliosis |

To determine the robustness of our guiding criteria (Fig. 2i), we asked 10 independent researchers with expertise in either human or rodent ripple detection to review candidate ripples from 9 patients (100 events x 18 channels; Fig. 3b; Supplementary Fig. 3). The agreement ratio varied widely for ambiguous events, but events meeting our criteria were widely accepted. Definite non-ripples were rejected by most raters (Supplementary Fig. 3b). For further analysis, we proceeded with fine-tuning the ripple detection parameters and using our criteria for ripple, IED and false positive classification.

### SPW-Rs are localized to the CA1 and subiculum in the human hippocampus

In our mice recordings, SPW-Rs were confined to the CA1 subfield of the hippocampus and have been reported in the subiculum in other studies^19,54^. To identify the spatial features of SPW-Rs, we localized human macrocontacts and microwire bundles to hippocampal subfields with the Automatic Segmentation of Hippocampal Subfields (ASHS) segmentation pipeline^55^ (Fig. 3a, Table 1). While the microwire bundle at the head of the depth electrode could be localized on the CT-scan, the precise locations of individual microwires could not be determined due to limited resolution (Fig. 2e, Supplementary Fig.4). In total, 20 macrocontacts and 19 microwire bundles inside the hippocampal formation were included for ripple analysis (Table 1). Most macrocontacts were in CA1 (n=15), with others in DG (n=2), subiculum (n=1) or white matter (n=2, with corresponding microwires in the hippocampus) (Supplementary Fig. 4). Microwire bundles were distributed across DG (n=6), CA1 (n=2), and subiculum (n=5). There were 6 microwire bundles that recorded across DG and neighboring subregions.

Our rodent results suggested that ripple frequency and power decrease with distance from the CA1 pyramidal layer. Therefore, we hypothesized that events with high power with narrowband high frequency identify human ripple-positive channels. To test this, we computed peri-event power spectral density (PSD) around automatically detected candidate ripples, as described above (±100 ms) and corrected the 1/f aperiodic component of the peri-event PSD (Fig. 3c; Supplementary Fig. 6). Macroelectrodes showed a bimodal distribution of peak frequencies: below or above 60 Hz (Fig. 3d). Manual review revealed that events detected on channels with ≤60 Hz peaks were all false positives, except for one channel in CA1 showed 11% of putative ripples (Supplementary Fig. 7). In contrast, about half of the channels with peaks above 60 Hz contained verified ripples. This metric captured most ripple-positive channels but failed to exclude 50% of ripple-negative channels (Supplementary Fig. 8). Thus, peri-event PSD can reliably exclude ripple-negative channels. For microwire bundles, one wire per bundle was selected based on the presence of a ripple peak in the PSDs (Fig. 3e,f). Of the 19 bundles, 11 contained at least one channel with a ripple peak. If multiple channels showed a ripple peak, the one with the largest peak power was selected for further analysis. Five of these channels showed manually verified ripples (Fig. 3f; Supplementary Fig. 9), while bundles lacking >60 Hz peaks showed no ripples upon visual inspection, confirming our method’s reliability.

Human ripples were primarily found in CA1 and subiculum, and were absent from DG electrodes, aligning with rodent findings (Fig. 3g). In contrast, IEDs were detected across all subfields. Approximately 65% of CA1/subiculum channels lacked ripples, likely due to their distance from the pyramidal layer or tissue pathology. These findings emphasize the importance of combining anatomical and physiological criteria to select optimal CA1/subiculum channels for ripple detection.

### Semi-automated separation of ripples and false positives in humans

Even after selecting optimal channels located within CA1 and subiculum and exhibiting >60 Hz peak, true ripples had to be separated from false positives, including IEDs and other HFOs. Visual inspection of all automatically detected ripples revealed high false positive rates (13% to 87%) across both macrocontacts and microwires (Fig. 4 a-c). False positives were mainly HFOs, which did not meet the criteria for 3 cycles. Pre-curation ripple rates during sleep were 6.8/min on macrocontacts and 6.3/min on microwires. After manual exclusion of false positives, these rates dropped to 3.1/min and 2.7/min, respectively, highlighting the need to control for false-positives.

**Fig. 4.**
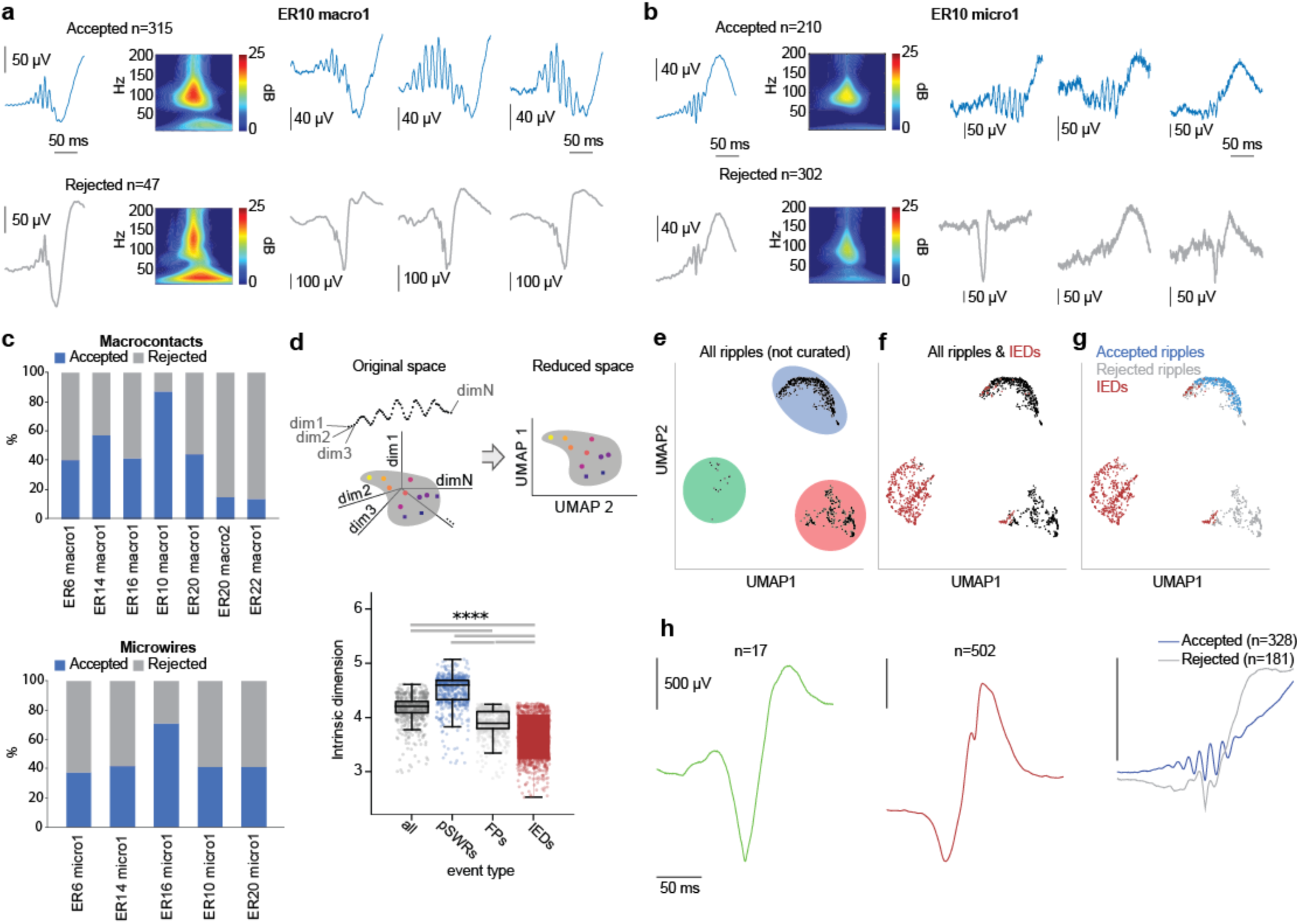
Manual curation of detected ripples reveals a significant proportion of false positives. **a** Average peri-ripple LFPs, wavelet spectrogram, and examples of accepted and rejected events from an example macro-contact. **b** Average peri-ripple LFPs, wavelet spectrogram, and example events of accepted and rejected events of an example microwire channel. **c** Proportion of accepted and rejected events across macro-contacts and microwires based on visual inspection. **d** Schematic of UMAP dimensionality reduction (top) and Comparison of the intrinsic dimension across different categories of events, adapted from^57^. **e** UMAP projection of all automatically detected ripples, before manual curation (n_neighbors=100, min_dist=0). The events are projected in a common embedding with IEDs detected in the same channel (not shown). Colored circles represent four distinct clusters of events**. f** Same as (e) but including IEDs. Note that some events project together with IEDs, potentially allowing their separation by UMAP before manual curation. **g** Same as (f) but after manual curation of automatically detected ripples. Blue: accepted, Grey: rejected. **h** Average waveform of three clusters in (e). The green and red colors correspond to the colors of the clusters in (e). The blue cluster was manually curated, demonstrating the mixture of accepted (blue) and rejected events (gray) in this cluster. Note that the rest of the clusters near IEDs show IED-like waveforms.

Ripples detected on macrocontacts and microwires showed similar peak frequencies (macro: 86±0.2 Hz, micro: 90±0.3 Hz; mean±SEM; Cohen’s d=0.26) and durations (macro: 64±0.39 ms vs. micro: 61±0.5 ms; Cohen’s d=0.16). However, macrocontact ripples showed larger amplitudes (macro: 1.34±0.004 vs. micro: 0.94±0.006; Cohen’s d =1.21; Supplementary Fig. 10). Most putative ripples showed peak frequencies below 150 Hz, below the frequency band associated with pathological HFOs^2,26^. Interestingly, hippocampal ripples in one frontal lobe epilepsy patient showed waveforms resembling rodent SPW-Rs from the pyramidal cell layer with peak frequency ∼140 Hz (Supplementary Fig. 10c).

False positives occurred across all frequency bands, including the range of putative ripples (80-250 Hz). This is possible because PSDs do not depend on the number of cycles we used to define ripples. For instance, events with fewer than three cycles in a specific frequency band can still produce a spectral peak on the PSD. False-positives were shorter in duration (macro: 64.64±0.39 vs. 55.45±0.29; Cohen’s d =0.39; micro: 60.75±0.53 ms vs. 51.11±0.38; Cohen’s d =0.44) and smaller in amplitude (macro: 1.34±0.004 vs. 0.86±0.005, Cohen’s d =1.18; micro: 0.94±0.006 vs. 0.64±0.005, Cohen’s d =1.01) than putative ripples (Supplementary Fig. 10).

Previous studies have identified IEDs as the primary source for false positives in ripples detection^22,31,35^ and our analysis confirmed this. Despite discarding IEDs before ripple detection through an automatic detector, some missed IEDs were falsely detected as ripples. Due to their distinct sharp morphology, IEDs were easy to differentiate from ripples by visual inspection (Fig. 4a,b).

HFOs were more challenging to separate from putative ripples. To separate events based on morphology, we applied uniform manifold approximation and projection (UMAP^56^) to ripple and IED snippets (Fig. 4d; see Methods). This approach draws on the fact that events that are similar will be close in the high-dimensional feature space, where each dimension represents one timestamp of the snippet. By using UMAP, we can reduce the dimensionality of the cloud of events while keeping the topological properties of the original space (Supplementary Fig. 11a,b). This strategy has already been successfully applied to rodent SPW-Rs, demonstrating its effectiveness to capture the variability of their features^57^. We confirmed that the intrinsic dimension of the events was also four for human events^57^. Interestingly, the intrinsic dimension of manually identified ripples was significantly higher than that of IEDs (Fig. 4d). We evaluated how well UMAP could separate all candidate events (including SPW-Rs, HFOs, and IEDs) clustered in both the original and reduced space, using several clustering validation metrics. Ripple-IED segregation depended on the parameters used to make the snippets (e.g. window size, sampling frequency, etc.) in a consistent manner between the original and 4D space (Supplementary Fig. 11d)^58,59^. The parameters that optimized event segregation were ±10ms duration, no extra channels, 400-500 Hz lowpass filtering, detrending and no z-scoring (Supplementary Fig. 12). Sampling frequency did not have a significant effect (Supplementary Fig. 12).

We used these parameters to perform the UMAP-aided semi-automatic curation. For this, we developed ‘ripmap’, a user-friendly open-source toolbox that allows versatile inspection and curation of events (https://github.com/acnavasolive/ripmap; see Methods). We found that IEDs falsely detected as ripples often clustered together with IED events and formed a separate cluster from putative ripples (Fig. 4e,f; see Supplementary 13 for UMAP embedding of all ripple-positive channels). This suggested that UMAP can effectively remove falsely labeled IEDs from ripples, reducing the need for manual curation by up to 45%. However, non-IED false positives lacked consistent features and did not show clear oscillatory patterns (defined by having ≥3 cycles) (Fig. 4g,h). These false positive events did not cluster with IEDs and often overlapped with putative ripples, making them harder to automatically separate (Fig. 4g). Overall, UMAP is useful for separating IEDs and HFOs from SPW-Rs, but not sufficient. Expert curation is still needed to separate SPW-Rs.

### Relationship between SPW-Rs on macrocontacts and microwires

A key question in the study of human memory is whether macrocontacts, which sample tens of thousands of neurons, can detect localized events such as ripples. Macro-micro hybrid electrodes enabled us to directly relate ripple detections from macro- and micro-contacts. Ripples detected on five microwire bundles were also detected on macrocontacts, if they were located in the same subfield or nearby white matter. We examined the microwire recordings during “macro-ripples” (ripples detected on macrocontacts). Extracting microwire signals from a ±100 ms window around macro-ripple peaks, we computed PSDs and counted the number of microwires with a peak above 60 Hz (Fig. 5a-b, f-g). Most accepted macro-ripples were coupled with simultaneous ripples on microwires, whereas false-positives on macrocontacts showed little or no corresponding microwire ripples (Fig. 5c,h; Supplementary Fig. 14). The peak frequency and amplitude of these macro-ripples were consistent regardless of the number of microwires involved (Fig. 5d-e, i-j). Two CA1 macrocontacts with ripples lacked microwire correlates, as, the microwires in these patients were located in different subfields-DG and subiculum, respectively. In summary, ripples detected on macrocontacts are also seen on microwires and vice versa when all channels are in the CA1 or subiculum.

**Fig. 5.**
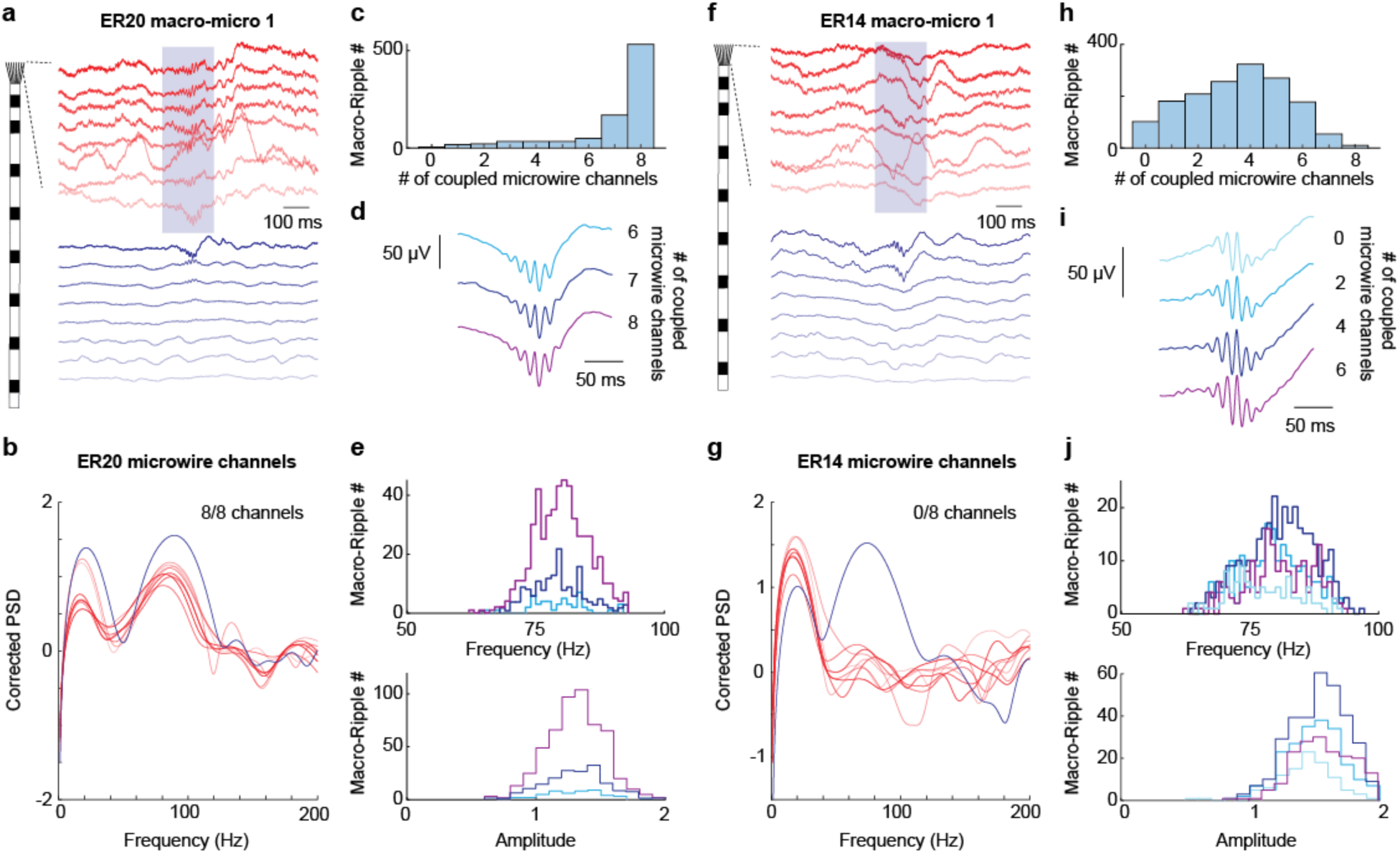
Coupling between ripples on macrocontacts and microwires. **a** Example ripple detected on a macrocontact (blue) coupled with microwire channels (red). **b** The number of coupled microwire channels were estimated based on peri-macro-ripple PSD of the microwire signals. In this example, all eight microwire channels show a ripple peak. **c** Distribution of the number of coupled microwire channels of the electrode in (a). The colors correspond to the channels in (a). **d** Average peri-macro ripple LFPs as a function of the number of coupled microwire channels, indicated by the number next to the waveforms. **e** Ripple peak frequency and amplitude distribution as a function of the number of coupled microwire channels, color coded as in (d). **f** Same as (a) but in this example, none of the microwire channels showed a ripple peak during a macro-ripple event. **g** Same as (b) but for the example in (f). **h** Distribution of the number of coupled microwire channels of the macro-micro pair in (f). **i** Same as (d) but for the electrode in (f). **j** Same as (e) but for the electrode in (f).

### Single unit responses to IEDs and ripples

Human ripples can be detected on macro-contacts, whereas neuronal spiking is detected in microwires that may be in a different subfield. To estimate peri-ripple spiking modulation across all hippocampal subfields, we first identified putative principal cells and interneurons by their spiking frequency and profile (see Methods) in APP/PS1 mice (314 principal cells, 167 interneurons) and in epilepsy patients (102 principal cells, 73 interneurons, Fig. 6). As expected, mouse SPW-Rs were associated with a brief burst of firing of principal cells and interneurons in CA1, CA2, CA3^16,17,42^ and DG (Fig. 6a). The increase in neuronal firing rates was more widespread than the observed LFP ripple oscillation (Fig. 2).

**Fig. 6.**
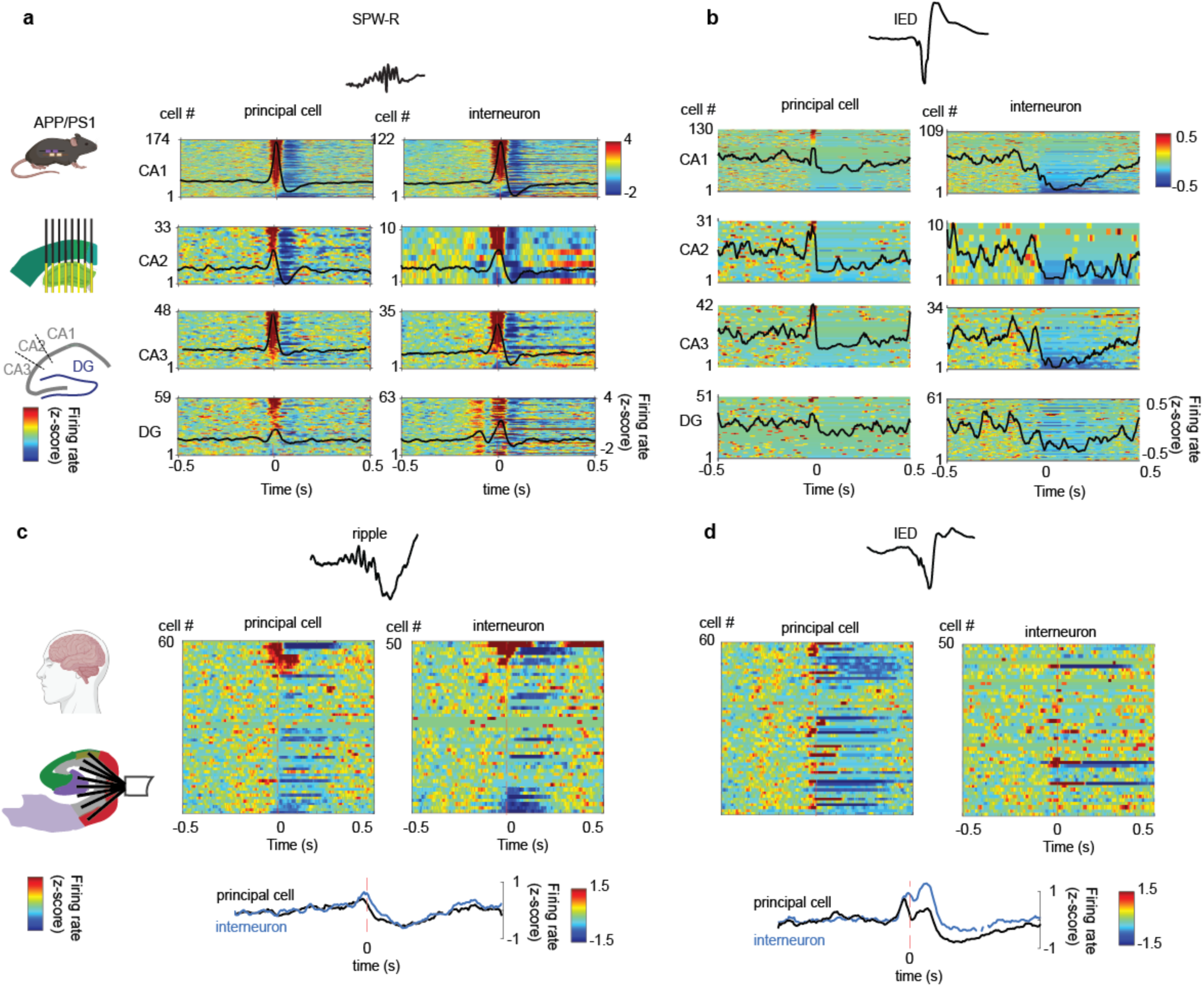
SPW-Rs/ripples and IEDs show distinct modulation patterns of neural spiking in rodents and humans: **a** Z-scored peri-SPW-R spiking in all recorded putative principal cells and interneurons from APP/PS1 mice. Neurons are sorted by hippocampal subfields and by the strength of peri-SPW-R positive neuronal modulation. Black traces, mean z-scored firing rates of all cells around SPW-Rs. Color axis: z-scored firing rates [−2 to 4]. **b** Same for peri-IED spiking. Neurons are sorted by hippocampal subfield and by the strength of peri-IED positive neuronal modulation. Color axis and black traces represent mean z-scored firing rates [−0.5 to 0.5]. **c** Top: Z-scored peri-ripples spiking in putative principal cells and interneurons recorded from epilepsy patients. For patients with multiple microwire bundles, only neurons recorded from microwires within the same depth electrode used for ripple detection are shown. Neurons are sorted by strength of peri-ripple positive neuronal modulation. Bottom: mean z-scored firing rates of all cells around ripples. Color axis: z-scored firing rates [−1.5 to 1.5]. **d** Same for peri-IED spiking. Only IEDs detected on the same ripple+ channels as in (**a)** were analyzed. Neurons are sorted as in **c** to compare directly the peri-IED and peri-ripple modulation for each neuron. Bottom: - mean z-scored firing rates of all cells around IEDs [−1.5 to 1.5].

We also analyzed the neuronal modulation by IEDs (Fig. 6b). The dominant effect of IEDs was a prolonged depression in firing rates of both principal cells and interneurons. Spike depression lasted approximately 0.5 seconds and was more pronounced in CA1, CA2 and CA3 compared to DG (Fig. 6b). As IEDs represent hypersynchronous population bursts, it appeared counterintuitive that IEDs caused only mild increases in neuronal firing rates at onset. We attribute this to spike-detection failure caused by superimposition of multiple units during strong synchrony.

Next, we evaluated the spatial distribution of human ripple modulation. A strong positive modulation was seen in neurons recorded on the depth electrode used for ripple-detection (Fig. 6c). Firing rates of neurons on distant depth electrodes in the same patient did not show modulation (Supplementary Fig. 15a). Neurons modulated by IEDs differed from neurons modulated by ripples, indicating the validity of our LFP pattern separation method. Overall, IED modulation of principal cells was similar to APP/S1 mice, with prolonged suppression in firing rates (Fig. 6d). This effect was less prominent in interneurons and pyramidal neurons on distant depth electrodes from the same patient (Supplementary Fig. 15b). The firing rates during false-positives were more variable possibly due to a mix of no modulation and IED-like modulation (Supplementary Fig. 15 c-d).

## Discussion

With the introduction of high-resolution electrodes and advanced computational methods, the neuroscience of human memory has entered a new era. Recently, there have been numerous publications relating “ripple activity” to memory processes. However, the field is limited by variable ripple definitions, low spatial resolution and imprecise localization of clinical macroelectrodes, and arbitrary detection thresholds^29^.

Our work identified hippocampal ripples in humans based on key spatial, spectral, and morphological features of SPW-Rs and IEDs obtained from high-resolution recordings in mouse hippocampus. Human ripples and IEDs matched rodent SPW-Rs and IEDs at multiple spatial scales (LFP and unit level).

### Key features of SPW-Rs and IEDs in rodents and humans

SPW-R features in mice included (1) tight ripple localization to the CA1 pyramidal layer, (2) a narrow-band peak within 100-250 Hz, (3) multiple ripple cycles (≥3) in the raw LFP, (4) differing polarity of the sharp-wave component in the cell-body (positive) and dendritic (negative) CA1 layer, and (5) accompaniment with a burst of pyramidal and interneuron firing followed by quiescence. Human ripples were also highly localized to CA1/subiculum and absent in DG, and showed a narrow band frequency peak between 80 and 120 Hz with multiple LFP cycles. Human pyramidal cells and interneurons demonstrated a burst of activity followed by silence, analogous to rodent activity, but to a lesser degree (likely due to sparser sampling). Conversely, rodent IEDs had (1) a broad spatial field, (2) higher voltage (∼5-15 fold), (3) a broadband peak, and (4) a decrease in pyramidal and interneuron firing after IEDs. Human IEDs had similar features, except fewer single neurons were modulated, possibly due to the varying distances of the individual microwires from the IED source. A previous study using rigid microwires showed consistent local activation and global suppression of human neuronal firing by IEDs^60^ similar to our rodent findings.

### Human hippocampal ripples can be detected on clinical macroelectrodes

While macrocontacts sample from tens of thousands of neurons^61,62^ they could detect ripples. Ripples were detected on both macrocontacts and microwires in CA1 and subiculum showing similar peak frequencies. However, electrodes residing ∼1.5 - 5 mm away from the CA1 pyramidal layer did not detect ripples. Most ripples detected on macrocontacts were also visible on multiple microwires, and vice versa. In two cases, a small percentage of ripples was detected on CA1 macroelectrodes, but not in microwires. We interpret this discrepancy by assuming that the microwires were located far from the CA1 pyramidal layer. In the neurosurgical transverse approach, where macrocontacts target the CA1, the microwires extend to more medially located structures, i.e. DG and subiculum, (Fig. 1d).

### Recommendations for ripple detection in human recordings

Based on our findings, we propose the following sequence of steps to isolate ripples from IEDs and other false positive events in the human hippocampus (Fig. 7).

***1. Localization of electrodes to hippocampal subfields.*** Given the highly localized nature of SPW-Rs to CA1 and subiculum, localizing electrodes is critical. We co-registered the post-operative cranial CT to the pre-operative MRI scans, used the ASHS segmentation pipeline to identify hippocampal subfields, and confirmed the electrode positions by visual inspection (Supplementary Fig.4). We recommend discarding electrodes in DG, because the false positive rate is expected to be high.
***2. IED detection and cleaning.*** Next, we recommend removing IEDs from the raw LFP signal prior to ripple detection. This is important because IEDs falsely raise the baseline threshold for detection of other high-frequency transients, including ripples. Moreover, IEDs can be falsely detected as ripple transients because of their wide-band power increase caused by the high-voltage sharp-transient (Supplementary Fig. 5).
***3. Discarding electrodes with PSD peaks <60 Hz.*** Next, we recommend excluding suboptimal channels by examining the average peri-event PSD. If the PSD of signals recorded during sleep lacks a high-frequency (<60 Hz) peak, most detected events will be false positives. By selecting channels with a narrowband peak above 60 Hz, we eliminated 49% of the detected events (from 57,276 to 29,278 events).
***4. Visual and morphological verification of candidate ripple events.*** Because averaged LFP waveforms can be misleading (Fig. 4a,b), individual candidate events must be visually inspected. UMAP clustering provides an unbiased approach to group events by LFP similarity. The toolbox we developed for this, “ripmap” was helpful in separating IEDs from ripples and increased the efficiency of our selection, by discarding up to 45 % of false-positives.

**Fig. 7.**
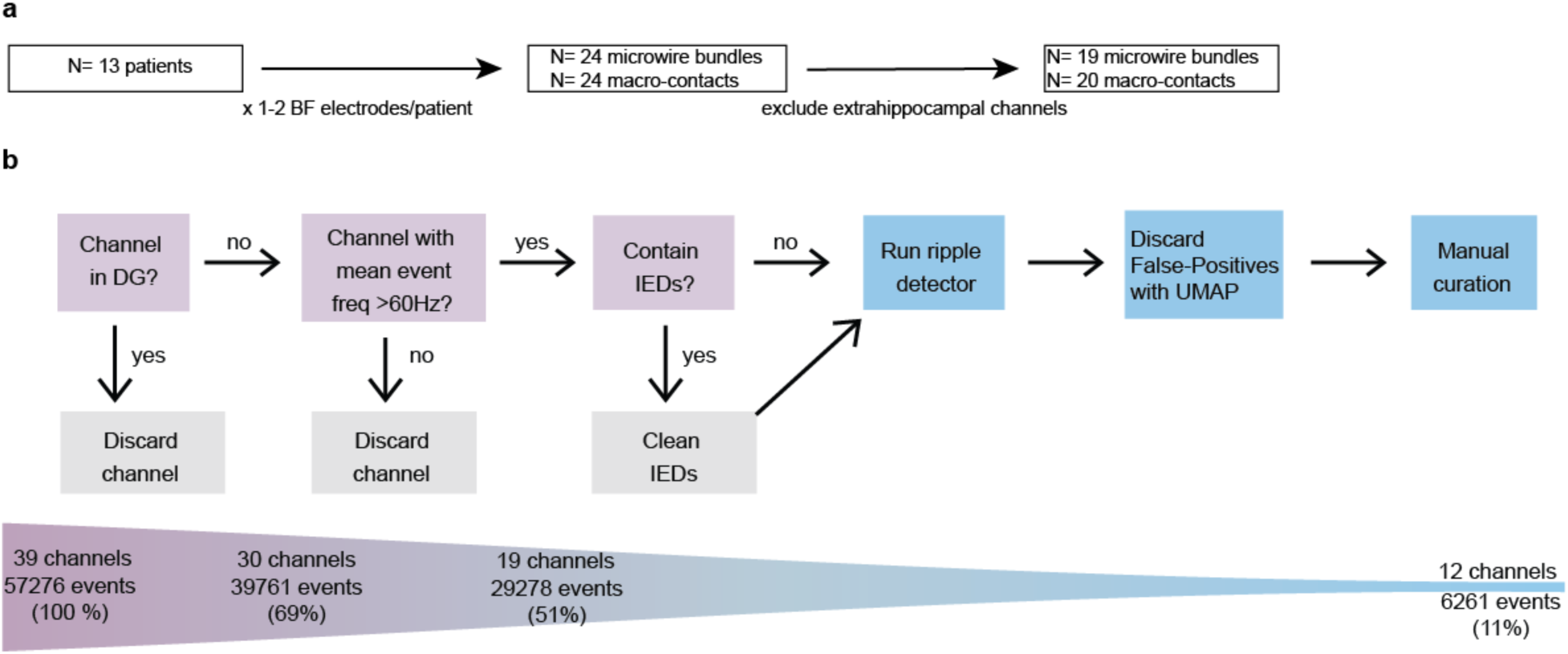
Human ripple detection pipeline. **a**. Channel inclusion and exclusion in this study. In total, 41 channels were used. **b**. Ripple detection pipeline and the number of channels events after each step.

Overall, only 11% of automatically detected candidate ripples in our dataset were confirmed as physiological ripples (n=6261 out of n=57276 events). We acknowledge that our approach prioritizes reducing false-positive rates over maximizing true-positive detection. Our ripple event rate was around 3 events per minute, which is near the lower limit compared to other reported rates ranging from 0.35 to 30 per minute^22,26,29,63^ and lower than in rodents^37,50^. This difference may be partially explained by our strict curation criteria. Additionally, IEDs in epileptic patients may suppress ripples during sleep, as shown in rodents and awake patients^25,37,50^.

### Awake ripples

In this study, we focused on ripples during sleep. Identifying ripples during awake states is further challenged by greater muscle artifacts and fluctuating arousal states, and contamination with other physiological events with power in the high-frequency band such as gamma oscillations.

Most of the human studies on ripples have been performed during waking cognitive tasks, reporting a relationship between ripple occurrence and successful encoding and retrieval of autobiographical, verbal, and visual information^33–35^. Given the predominance of true SPW-Rs in low-cholinergic states in rodents^6,64^, observing ripple activity during effortful states is a conundrum. Theta coupled with gamma, not SPW-Rs, are observed in awake rodents during ambulation^65^. However, during rapid changes between immobility and ambulation, SPW-Rs may preconsciously “prime” hippocampal circuits and assist with both route planning and memory selection in rodents^66–68^. It is possible that rapid changes in brain states may occur in primates during pre-attentive and attentive states. Alternatively, it is also possible that high-frequency events deemed as “ripples” in some human studies are actually transient gamma patterns, which are promoted by attention and increased cholinergic tone^6,69^. Future studies combining hippocampal recordings with measurements of arousal, such as pupil diameter, may disambiguate ripples from gamma oscillations during awake states. In addition, development of feature-based detection methods would allow both more specialized oscillatory pattern detection and greater independence from noise thresholds^70^.

### Limitations

We acknowledge some spatial uncertainty in our findings in the human brain. Ideally, definitive subfield location could be determined by examining histological sections of the resected hippocampus, which was not available to us. Co-registration of pre- and post-operative imaging to idealized subfield segmentation and subtle brain shift may introduce uncertainty to electrode localization. A recommended improvement is to cut the microwires at predetermined distances, so that a laminar profile of the LFP can also be recorded in humans. This will allow identification of ripples and their evoking sharp wave component.

### Future Directions

Our rigorous, cross-species approach aims to benefit the human cognitive neuroscience community and accelerate translational research on memory function. Moreover, our findings will be useful to the epilepsy and Alzheimer’s communities, with growing awareness of IEDs’ pathological role in memory disruption and disease progression^25,48,71,72^. Reliable methods to distinguish hippocampal ripples from IEDs and other high frequency oscillations in epileptic brain tissue are crucial to mechanistic understanding of brain-behavior dynamics during memory tasks. In the future, neuromodulation techniques via closed-loop IED suppression or selective ripple enhancement^4^ may offer a promising means of memory restoration in clinical populations.

## Acknowledgements

We thank Karl Rössler and Sebastian Brandner for the human SEEG implantations; Katja Kobow for providing the histopathological findings of the patients; Jay Jeschke for help with human electrode localization; Esha Brahmbhatt and Deren Aykan for help with animal habituation; Mursel Karadas for the rodent treadmill design; Nicholas Paleologos, Noam Nitzan, Michael D Hadler and Samuel McKenzie for rating events in the human ripple survey; Thomas Hainmüller for feedback on the manuscript.

We would like to acknowledge Corticale SRL (Genoa,Italy) for providing the SiNAPS probes, and NeuroNexus (Ann Arbor, MI) for their contribution of the data acquisition system and Radiens software. We further acknowledge both Corticale and NeuroNexus for training and support making this research possible.

This research was supported by:

DFG Fellowship MA 10301/1-1 (AM), NYU FACES (AM)

NOMIS fellowship (AN-O)

R01 NS127954 (AL), K23-NS 104252 (AL)

NYU Department of Neurology

NIH MH122391 (GB)

U19 NS107616 (GB)

## Author contributions

AM, JS, AN-O, GB and AL designed the study. AM, MV, HH and AD collected data. AM, JS, AN-O and SH analyzed the data. AN-O developed the “ripmap” toolbox. AL and GB supervised the project. AM, JS, GB and AL wrote the manuscript.

## Methods

### Rodent Neurophysiology

#### Rodents

We used male APP/PS1 hemizygous mice (n = 5, age 3-9 months, JAX Stock No. 004462), These mice express mutations of two genes associated with Alzheimer’s disease (AD): amyloid precursor protein (APP) and presenilin-1 (PSEN1) and demonstrate neuronal hyperexcitability and alterations in cortical oscillations ^73,74^. They show early development of interictal epileptiform discharges (IEDs) ^37,38,53^, but seizures are very rare, allowing to study IEDs and physiologic events, without the influence of hippocampal sclerosis or postictal effects.

#### Rodent Handling and Surgery

All experiments were performed in accordance with the Institutional Animal Care and Use Committee at New York University Medical Center. Mice were housed at a vivarium on a 12-hour light/dark cycle and were handled daily and accommodated to the experimenter before the surgery and head-fixed recording. As described previously ^39^, for head-fixation, mice were implanted with a custom 3D-printed headpost^75^ (Form2 printer, FormLabs, Sommerville, MA), attached to the skull with dental cement (C&B Metabond, Parkell, Edgewood, NY) under isoflurane anesthesia. A stainless-steel ground screw was implanted above the cerebellum. After recovery from surgery for at least 7 days, mice were gradually habituated to the head-fixation for at least 5 consecutive days. The head-fixation system was equipped with a treadmill (https://github.com/misiVoroslakos/3D_printed_designs/tree/main/Treadmill_Rinberg), allowing the animals to walk freely during training and recording sessions. After the habituation was complete, a craniotomy was performed above the hippocampus (craniotomy dimensions 2.5*0.5 mm, center coordinates 2 mm posterior to Bregma, 1.5 mm lateral to midline) the dura was removed, and the craniotomy was temporarily sealed with Kwik-Sil (World Precision Instruments, Sarasota, FL).

#### Rodent Data Acquisition

Acute recordings were performed on the day after the craniotomy to allow for the effects of isoflurane to fade. Recordings were performed using 8-shank, 1024-channel active CMOS-based probes (SiNAPS, NeuroNexus, Ann Arbor, MI; Corticale SRL, Genoa, Italy^39,40^, Fig. 3a). These probes provide simultaneous recordings from all 1024 electrodes, distributed over 8 shanks x 128 channels in a two-column configuration (inter-shank distance of 300 µm, center-to-center electrode pitch ∼30 μm, total recording area of 2100×1924 µm). The probe was gradually inserted to the target depth using a manual micromanipulator (MM-33, Sutter Instruments, Novato, CA). During insertion, live monitoring of the electrophysiological signal was used to establish that the desired depth had been reached. The data was digitized at 20 kS/s using Smartbox Pro and Activus SiNAPS interface box and visualized using RadiensTM Allego software (NeuroNexus Technologies, Ann Arbor, MI). At least 20 minutes were allowed for equilibration before data collection, and 2-hour sessions were recorded per mouse. After each session, the craniotomy was resealed with Kwik-Sil, and mice were returned to their home cages. For some mice, analogous craniotomy and recording were also performed on the contralateral hemisphere.

#### Data Analysis

All analyses were executed with custom written scripts in Matlab 2022a (MathWorks). Scripts were adapted from https://github.com/buzsakilab/buzcode and https://github.com/valegarman/HippoCookBook).

#### Rodent Channel localization

Identification of hippocampal subfields, and cellular layers, was based on established electrophysiological markers as described in ^39^. **CA1:** We automatically detected SPW-Rs on a single electrode from the CA1 pyramidal layer selected by the and calculated current-source density (CSD) profiles^76^ to delineate the stratum radiatum and pyramidal layer of the CA1/CA2 axis (Fig. 2b). For further analysis we chose the CA1 pyramidal channel with the highest high-frequency power (100-500 Hz) and the CA1 radiatum channel on the same shank with the largest negative wave during detected SPW-Rs. **DG:** Analogously, we computed CSD profiles of dentate spikes detected in the dentate gyrus^77^. Dentate spikes are brief large-amplitude LFP waves in the DG representing the firing of DG neurons in response to inputs from the entorhinal cortex. Dentate spikes show polarity reversal above the granule cell layer^77,78^. The border between current sinks and sources of SPW-Rs and dendritic layers^39^. **CA2 and CA3:** Identified neurons helped to delineate the pyramidal layers of CA3, CA2, and CA1. Separation of CA2 was performed based on peri-ripple firing histograms^39,42^.

#### Rodent IED detection

IED detection was performed on a CA1 radiatum channel (see above). The wide-band LFP signal (1250 Hz) was band-pass filtered between 20 and 80 Hz^37,50^ and transformed to a normalized squared signal (NSS). IEDs were detected by thresholding the baseline at 5 SD and thresholding peaks at 20 SD of the NSS for 50 to 250 ms. All events were then visually inspected and manually curated. If necessary, the SD thresholds were adjusted for individual subjects. For further analysis, detected events were realigned to the trough of the unfiltered LFP.

#### Rodent SPW-R detection

SPW-R detection was performed as previously described. We picked the CA1 pyramidal channel with the highest high-frequency (100 −500 Hz) power. The wide-band LFP signal (1250 Hz) was band-pass filtered between 130 −200 Hz using a 3rd order Butterworth filter and transformed to a normalized squared signal (NSS). Periods of IEDs (± 1s) were excluded. SPW-Rs were detected by thresholding the baseline at 2SD and thresholding peaks at 5 SD of the IED-free baseline signal. Events shorter than 30 ms or longer than 250 ms. The correct detection was confirmed by manual curation.

#### Frequency analysis

Event-triggered wavelet spectrograms were computed for the frequency range of 1-400 Hz using Morlet wavelet. Power spectral densities were computed using Welch’s average.

### Human Neurophysiology

#### Human Subjects

We conducted a retrospective analysis of intracranial recordings from 14 patients **(Table 1)** with drug-resistant epilepsy who underwent diagnostic stereotactic electrode implantation for identification of epileptogenic cortex. Patients in the preoperative epilepsy program of the Level 4 Epilepsy Center of Erlangen University Hospital, Germany, were recruited and consented from 2019 – 2022 years. The decision for implantation was made by the interdisciplinary epilepsy board. Ethical approval was granted by the “Ethik Kommission” of Friedrich-Alexander Universität Erlangen-Nürnberg (142_12B), and all patients consented to implantation of hippocampal macro/microwire bundles (Behnke-Fried Electrodes) in addition to standard iEEG depth electrodes (AdTech Medical, USA) in other temporal and extratemporal regions. One patient (ER11) was excluded due to insufficient anatomical information. Informed written consent for the implantation of 1 to 2 hippocampal electrodes with microwire bundles.

#### Electrode reconstruction and hippocampal subfield segmentation

Putative electrode localization was performed by using automated processes and expert review. The location of macrocontacts and microwires was determined by co-registering preoperative T1-weighted structural MRI and postoperative computed tomography (CT) with visible electrodes artifacts using FSL *flirt* affine registration. For descriptive purposes (Fig. 1e,Supplementary Fig. 4), we aligned each subject’s CT images to MNI152 T1-weighted template using CT2MNI152 ^79^. MNI coordinates for each macrocontact and microelectrode were extracted manually from the normalized CT overlaid on the MNI152 T1 MRI using the FSLeyes software. To localize individual microwire bundles to hippocampal subfields (e.g., CA1, CA2/3, DG), we used the Automatic Segmentation of Hippocampal Subfields (ASHS) toolbox^55^.

#### Human sleep scoring using iEEG signals

To identify sleep periods in human iEEG signals, we computed the delta to gamma power ratio on all macro-contacts and microwires^37^. Initially, all iEEG signals were down-sampled to 64 Hz and divided into 30-second epochs before calculating the spectrogram and power spectral density (PSD). For each iEEG channel, we derived the spectrogram and delta-to-gamma power ratio. The spectrogram was obtained using a continuous wavelet transform with a Morlet Wavelet, set to 10 voices per octave and a time-bandwidth product of 90, using MATLAB’s “wt” and “cwtfilterbank” functions. We focused on frequencies between 0.5 Hz and 30 Hz. PSD was computed through fast Fourier transform (FFT). Relative delta (0.5 to 4 Hz) and gamma (20 to 30 Hz) power were calculated by normalizing the power within each frequency band by the total PSD power. We plotted the spectrogram showing median power across all channels for each frequency band, overlaid with the median delta-to-gamma ratio across the entire recording. This visualization allowed us to set a subject-specific delta-to-gamma threshold to distinguish wake and sleep epochs. Epochs with a delta-to-gamma ratio exceeding this threshold were classified as sleep, while those lasting less than 5 minutes were considered wake epochs and excluded from analysis.

#### Human macroelectrode and microwire recording

In addition to standard depth macroelectrodes (Ad-Tech Medical), patients were implanted with 1 to 2 macro/microwire hybrid electrodes (Behnke-Fried electrodes, Ad-Tech Medical) in mesial temporal lobe structures. Electrode trajectories were planned by the clinical team based on surface EEG, seizure semiology, and additional diagnostic modalities and without respect to this study. Microwire bundles consisted of eight high-impedance wires and one low-impedance wire, which served as reference. Prior to implantation, all microwires were shortened to extend 5 mm in front of the first macro-contact. Electrode positions were confirmed by an intraoperative MRI- and postoperative MRI- and CT-scan. Only recordings from temporal electrodes with hippocampal microwires were included in this study. Following surgery, patients were monitored continuously for 7 days upon withdrawal of antiepileptic drugs.

Data were acquired using an ATLAS recording system consisting of CHET-10-A preamplifiers and a Digital Lynx NX amplifier (Neuralynx Inc.). Signals were filtered with cutoff frequencies at 0.1 and 9,000 Hz and sampled at 2048 Hz for macroelectrodes and 32,768 Hz for microwires. The LFP signal from microwires was downsampled to 2,048 Hz. Line noise at 50 Hz, 100 Hz and 150 Hz was removed using a second-order infinite impulse response notch filter (irrnotch.m). Hippocampal macroelectrodes were re-referenced to the nearest white matter contact of the same electrode. The raw data were converted to int16 binary format for processing with buzcode analysis scripts in Matlab (MathWorks; https://github.com/ buzsakilab/buzcode).

#### Human Microwire channel selection

We selected one channel per microwire bundle to thoroughly inspect SPW-Rs. We extracted peri-event PSDs after automatically detecting putative SPW-R events after discarding periods around IEDs (± 1s) as described above but without any manual curation. Channels with less than 100 events were excluded. We then extracted spectral peaks from 1/f-corrected PSDs. If none of the channels showed a peak in ripple band, the bundle was considered SPW-R negative. If more than one channel showed a ripple band peak, the channel with the strongest peak amplitude was chosen for further analysis. In these channels, both IEDs and SPW-Rs were manually curated to reject false positive events.

#### IED detection in humans

IED detection was performed on the same channels used for SPW-R detection. The raw LFP signal was band-pass filtered between 20 and 80 Hz^37,50^ and transformed to a normalized squared signal (NSS). IEDs were defined by peaks beyond 10 SD of the NSS whenever the signal crossed 3 SDs of the NSS for 50 to 250 ms. All events were then visually inspected and manually curated. Whenever necessary, the SD thresholds were adjusted for individual subjects. For further analysis, detected events were realigned to the peak of the unfiltered LFP. For mice, events from behavior periods were discarded because of artifact contamination.

#### Human ripple detection

In humans, one microwire channel was selected based on visual inspection, and the nearest macroelectrode was used to detect SPW-Rs. Signals were band-pass filtered from 80 to 250 Hz using a 3rd order Butterworth filter. This wide ripple band was chosen to capture a diverse range of SPW-R events and to examine how electrode size and location affect peak frequency. Very few events had peak frequencies above 160 Hz, indicating that we were primarily detecting physiological SPW-Rs despite the wide frequency band. IEDs can exhibit large transient peaks of broadband power, which can falsely raise the detection threshold for SPW-Rs (Supplementary Fig. 4). To reduce the impact of IEDs on the average and standard deviation of the signal, one second of data around IED peaks was removed after rectifying the filtered signal. This process produced a ripple power time series, which was then z-score normalized. SPW-R detection was restricted to sleep epochs. Putative SPW-R events were defined by beginning and ending cutoffs exceeding 2 standard deviations and peak power exceeding 5 standard deviations. Additionally, events were required to last between 30 and 250 ms, and events separated by less than 30 ms were merged. All events were visually inspected and manually curated. One microwire bundle (located partially in the subiculum and/DG) with only 89 ripple events (0.3/min) were considered ripple-negative.

For recordings where macro-contacts on the same electrode as SPW-R+ microwire bundle, microwire snippets around the detected macro SPW-R events (e.g., if microwire shows ripple peak, how many channels show a ripple peak) were used the guide the manual curation in addition to the waveform.

#### Spectral peak extraction using FOOOF

Previous methods for extracting the peak frequency of candidate SPW-R events typically relied on filtered signals within a pre-defined ripple band (80-120 Hz in humans, 80-250 Hz in rodents). This approach often forced the identification of peaks within this band, even if there was no clear spectral peak in the power spectrum. To more accurately estimate the ripple peak frequency, we first extracted the power spectrum 200 ms around the ripple peak (±100 ms). The aperiodic (slope) component was removed using FOOOF^80^, resulting in a 1/f corrected power spectrum. Peaks were then identified from the corrected power spectrum using the findpeaks.m function, and those with an amplitude less than 0.2 (average PSDs) or 0.5 (individual PSDs) were discarded. If peaks were found between 30-180 Hz, the one with the highest amplitude was selected. If no peak was found in this range, the low-frequency peak (<30 Hz) was identified.

#### Spike sorting in rodents and humans

Spike sorting was performed semiautomatically using Kilosort 1 for human and Kilosort 2.5 for rodent recordings (https://github.com/MouseLand/ Kilosort), followed by manual curation of the waveform clusters with Phy2 (https://phy-contrib.readthedocs.io/). For human recordings spike sorting was performed separately for each microwire. Neurons were classified as putative pyramidal cells or interneurons using an automated cell type classification toolbox, CellExplorer^81^, as described previously^37^. CellExplorer separates interneurons and pyramidal cells according to trough-to-peak duration of their waveforms and autocorrelogram shape of their firing rates. Parameters are based on known distinctive features: briefly, hippocampal pyramidal cells typically have much wider waveforms than interneurons, are prone to bursting and their average firing rate is very low (< 2 Hz). By contrast, interneurons demonstrate higher firing rates and lack bursting behaviour^27,82,83^. Automatic classification was followed by manual verification.

#### Event inspection and curation with UMAP

Events detected with the SPW-R detector or IED detector can be further analyzed and curated by reducing their dimensionality and visualizing them in a 2D space. For this, we applied uniform manifold approximation and projection (UMAP)^56^ over all the detected events from both detectors. Before running UMAP with automatically detected events, we recommend aligning the negative or positive IED peaks (Supplementary Fig. 13a). To make the analysis versatile and user-friendly, we created an open-source toolbox (ripmap) that runs on python. ripmap first performs an analysis of the topological features of the data, by (i) computing the mean intrinsic dimension of the data in the original space, and (ii) characterizing the shape of the cloud using persistent homology. Then, data is reduced to a lower dimensional space and plotted in 2D. ripmap GUI offers the possibility to perform the dimensionality reduction with different parameters at the same time and interactively select the parameters that better segregate IED-detected from SWR-detected events. If events are clearly segregated, the user can visualize a clustering of the UMAP cloud by specifying “do_clusters = 1” in the GUI. Pressing “continue”, a new figure appears with a plot in the left of the UMAP cloud colored by the detector, and an interactive plot in the right of the same cloud colored by the cluster number, and the mean event of the cluster. The interactive plot allows to select any custom area and display the events within it. The aim is to easily relate the position of the UMAP cloud with the waveform of the events. However, if the UMAP cloud shows no segregation and clustering is not possible, an alternative curation approach can be taken. If the user specifies “do_clusters=0” in the GUI, the 2D UMAP cloud is projected into a 1D axis and binned in equidistant segments. The new figure shows the mean waveform of the events of each bin, which typically transitions from a SWR-like waveform to an IED-like waveform. Individual events can also be displayed.

##### Data processing and UMAP projection

UMAP input is a matrix of events × time, where events are both the detections from the IED and the SPW-R detectors, and the different time points the events represent a particular feature. This strategy has been previously applied in rodents to analyze SPW-R diversity^57^. The original space will have as many dimensions as samples has the event, and the coordinates of each dimension X will be the LFP value at sample X. This means that the form of the cloud in the original (and reduced) space will depend on how events are created or processed. The key parameters that we incorporated in the toolbox to generate the UMAP input snippets are win_size_umap, which specifies the window duration at each side of the center of the event; do_detrend, which sets if each event is detrended; and do_zscore, that sets if each event is z-scored. We explored how these parameters affected the topology of the SWR-IED cloud, in addition to three other ones: number of channels used (one vs two), sampling frequency (original, 2048Hz, vs downsampled, 1024Hz), and frequency of a low pass filter (200Hz, 300Hz, 400Hz, 500Hz or no filtering). The exploration was performed by computing two different cluster index metrics for all possible sets of parameters and all patients, and evaluate which set of parameters result on a better score (Supplementary Fig. 11, 12). The metrics were Density-Based Clustering Validation (DBCV) index^59^, and Silhouette^58^, taken from permetrics, ClusteringMetric^84^ and sklearn.metrics^85^, respectively. Metrics were computed for all possible sets of parameters. The extra channel that we used was the channel that better optimized the metric for each case. To check for potential topological biases, data labels were shuffled 100 times for every set of parameters, and the metrics were recomputed. We observed that, although small in most cases, there were some significant differences in the cluster index metric values for different parameters, so we analyzed data subtracting the mean metric value per parameter (Supplementary Fig. 12). This analysis allowed us to compute the optimal snippets to input UMAP. In order to create an UMAP embedding in ripmap, more parameters need to be specified. In our toolbox, the metric is always euclidean, but the number of components (n_components), the number of neighbors (n_neighbors) and the minimum distance (min_dist) can be set to any value. We have used n_components=4 to match the intrinsic dimension (see Supplementary Fig. 11), and a list of several n_neighbors and min_dist, with list_n_neighbors=[10, 50, 100, 200], and list_min_dists = [0.0, 0.1, 0.2, 0.3].

To create an embedding with UMAP, more parameters need to be specified. In our toolbox, the metric is always euclidean, but the number of components (n_components), the number of neighbors (n_neighbors) and the minimum distance (min_dist) can be set to any value. We have used n_components=4 to match the intrinsic dimension (Supplementary Fig. 10), and a list of several n_neighbors and min_dist, with list_n_neighbors=[10, 50, 100, 200], and list_min_dists = [0.0, 0.1, 0.2, 0.3].

##### Intrinsic dimension and persistent homology

The intrinsic dimension of the data in the original dimension was computed using angle-based intrinsic dimensionality (ABID) method^86^. ABID derives the theoretical distribution of angles among neighboring points and uses this to construct an estimator for intrinsic dimensionality. We compute the intrinsic dimensionality of the data using the same list of neighbors than for UMAP, list_n_neighbors=[10, 50, 100, 200]. Persistent homology diagrams^87^ were computed adapting the code from previously published works^88^. This method computes topological features of a space at different spatial resolutions.

Assuming that features that are persistent during a wider range of spatial scales will more likely represent true features of the space, the diagrams capture what topological features define the cloud in the original space. This analysis can be done at several levels (so-called “Betti numbers”), and each level describes a particular topological feature. For example, the first Betti number, β_0_, describes the number of connected components. Therefore, the number of persistent (long) bars in the diagram will reflect the number of unconnected components (or clusters)

##### Cluster suggestion and projection to 1D axis

If specified, clustering can be done over the 2D UMAP cloud. Clustering is done using hierarchical density-based spatial clustering of applications with noise (HDBSCAN)^89^. Note that clustering in this pipeline is meant to be used as a visualization and inspection aid, rather than an automatic clustering tool for SWR or IED type detection. The input parameters for HDBSCAN are 10% of the number of events for the min_cluster_size variable, and 0.05*min_cluster_size for min_samples. The interactive plot allows to draw any shape and visualize all the individual events inside that shape, allowing an instant evaluation of the performance of the event segregation, and an easy and fast curation. The selected events can also be saved to conform the final curated set. Alternatively, if the UMAP projection has no clear segregation and has the shape of a single cloud, we can explore the distribution of the events by dividing the cloud into equidistant sections. The number of bins into which it is divided can be specified by the parameter n_axis_bins. The mean waveform of all the events laying within that bin is then displayed, with the option to plot false positive (if labeled) separately using plot_fp_separately=True. The axis can be computed in two ways: from the pIEDs centroid to pSWR centroid (using axis_method=‘centroids’), or fitting all data into a quadratic line (with axis_method=’fit’, see topol_utils.fit_axis). There is an additional option that allows plotting the events over and below the axis separately (do_axis_grid=True). Both in the clustering and projection options, the mean waveform and individual events displayed can be set to have a different length to the one used in UMAP with the parameter win_size_show.

UMAP was done using the python library umap-learn (https://umap-learn.readthedocs.io/), signal processing tools were taken from scipy (https://docs.scipy.org/doc/; scipy.signal.detrend, scipy.stats.zscore). Clustering was done using HDBSCAN (https://pypi.org/project/hdbscan/). Tests were run on Python 3.11.2 with the following library versions: h5py==3.12.1, hdbscan==0.8.39, matplotlib==3.9.2, numba==0.60.0, numpy==2.0.2, permetrics==2.0.0, persim==0.3.7, ripser==0.6.10, scikit-learn==1.5.2, scipy==1.14.1, tqdm==4.67.0, umap-learn==0.5.7.

##### Data Visualization

Illustrations in Fig. 1a include images from Biorender (publication license was obtained). Statistical figures were plotted using Gramm Toolbox^90^.

## Code availability

’Ripmap’ is available on https://github.com/acnavasolive/ripmap, which also includes an example script for optimal SPW-R-channel detection in https://github.com/acnavasolive/ripmap/tree/main/select_channel.

