## Supplemental figures for "Spatiotemporal Patterns Differentiate Hippocampal Sharp-Wave Ripples from Interictal Epileptiform Discharges in Mice and Humans"

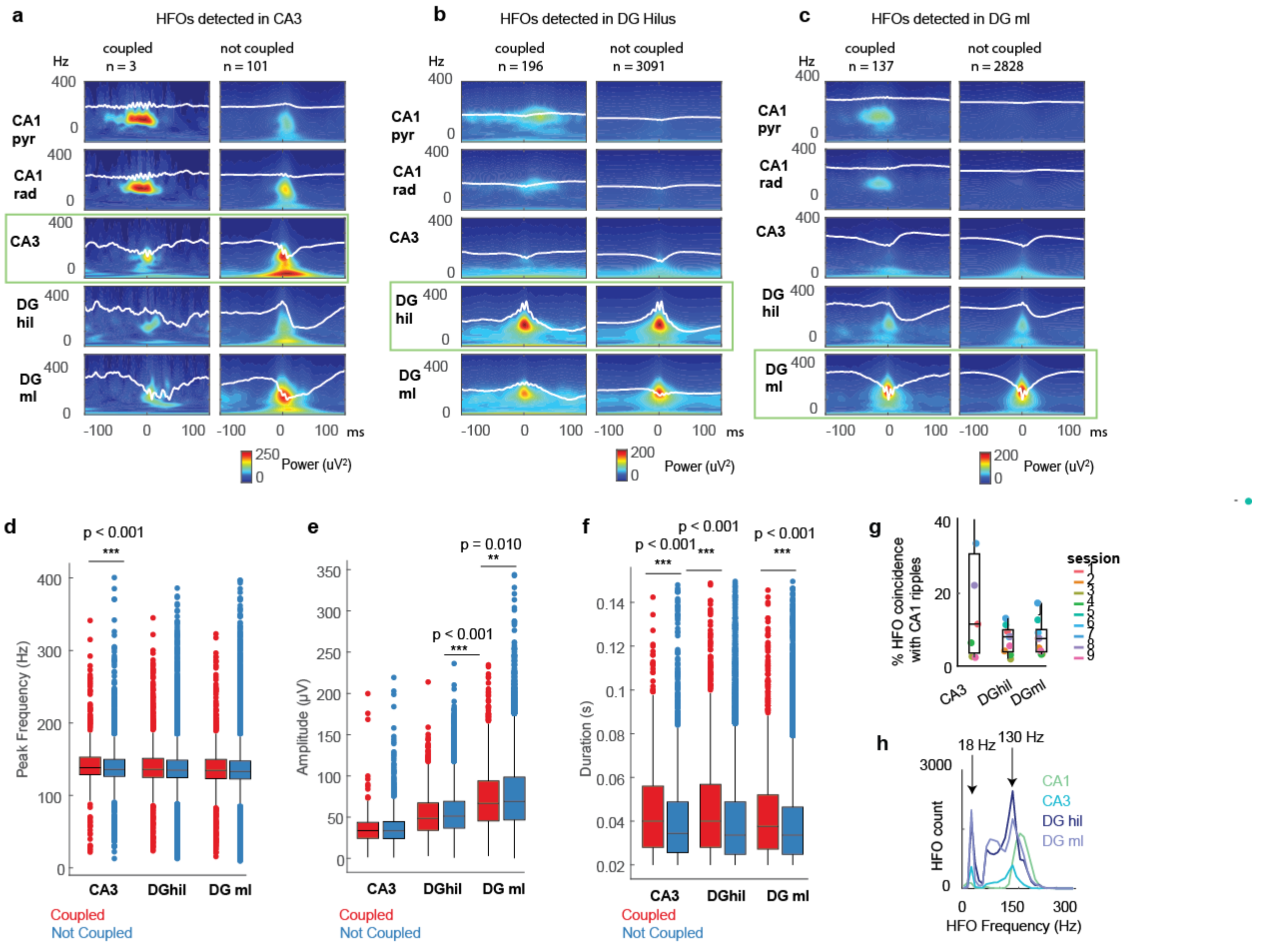


**Supplementary Fig. 1: Mouse high-frequency oscillations (HFOs) detected using SPW-R-detection parameters in CA3 and DG.** **a** Average LFP traces and wavelet spectrograms of HFOs detected on a CA3 detection channel from one recording session. The corresponding features computed on channels of different hippocampal subfields are shown: CA1 pyramidal layer (pyr), CA1 stratum radiatum (rad), CA3, DG hilus (hil) and DG molecular layer (ml). The detection channel is indicated by a green box. Left column: HFOs coupled to CA1-SPW-Rs (within ± 200 ms of the SPW-R peak in CA1). Right column: all other detected HFOs. **b** Same for HFOs detected in DG hilus. **c** same for HFOs detected in DG molecular layer. **d** Peak frequencies of HFOs detected in CA3, DG hilus and DG molecular layer and coupled to CA1-SPW-Rs (red box plots), compared to HFOs not-coupled to CA1-SPW-Rs (blue box plots). **e** Same for amplitudes of HFOs. **f** Same for HFO duration. **g**. Rates of coincidence of HFOs detected in CA3, DG hilus and DG molecular layer with CA1 SPW-Rs. Coincidence was defined as both events being detected within a 200 ms time frame. **h** Histogram of the peak frequencies of CA1 SPW-Rs and HFOs detected in different layers.

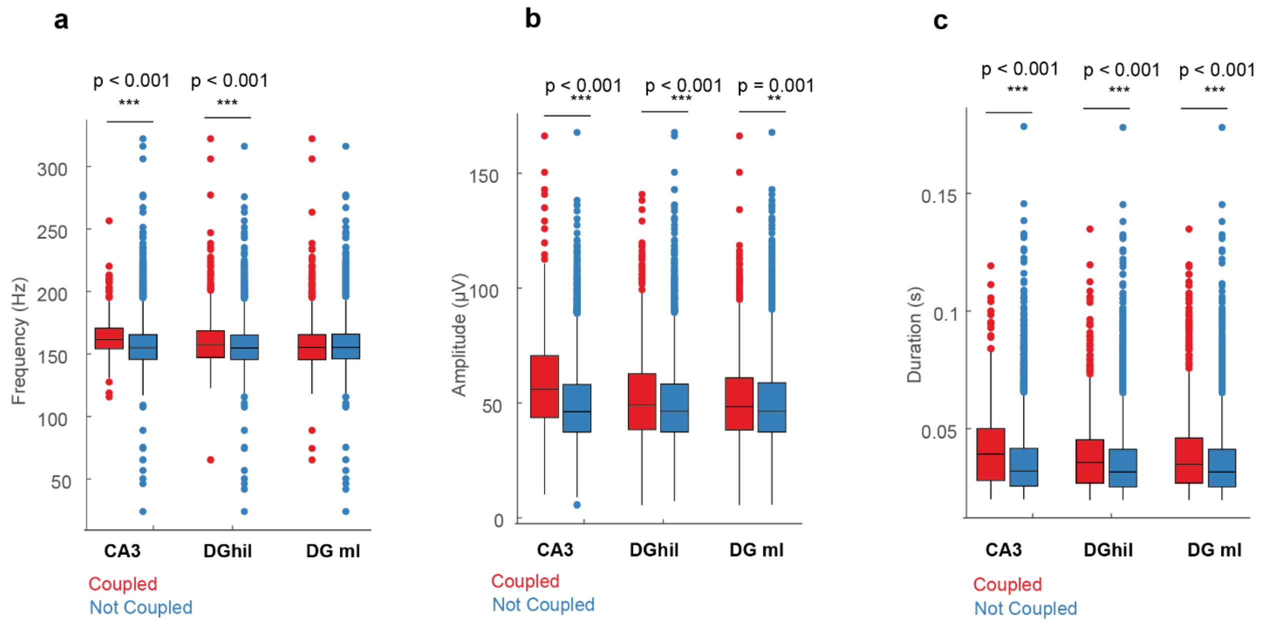


**Supplementary Fig. 2: Features of mouse CA1 SPW-Rs coupled to HFOs in other regions, compared to non-coupled SPW-Rs. a** Peak frequency of SPW-Rs detected in CA1, and coupled to HFOs detected in CA3, DG hilus, or DG molecular layer (red box plots), compared to peak frequencies of CA1-SPW-Rs not-coupled to HFOs (blue box plots). **b** Same for amplitudes of HFOs. **c** Same for HFO duration.

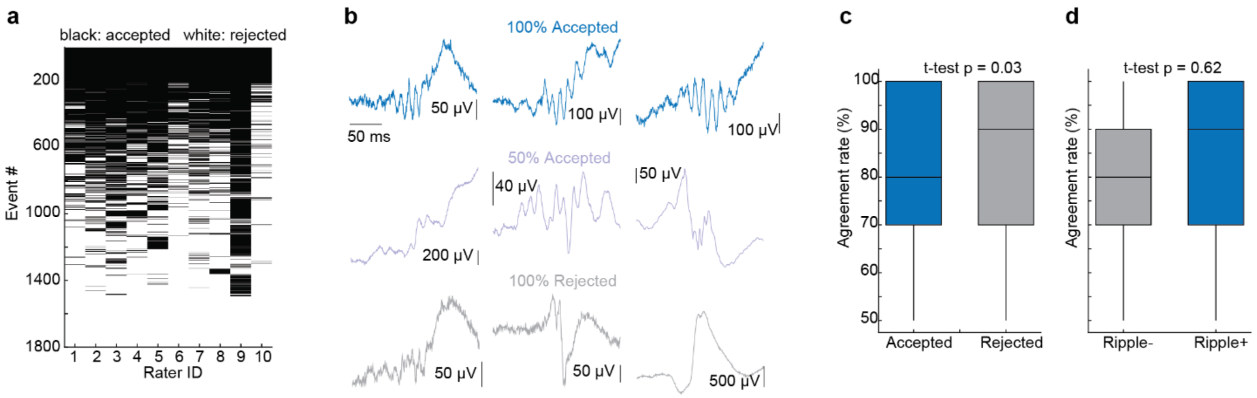


**Supplementary Fig. 3: Human ripple rating varies largely among individual evaluators. a** Rating from ten individual researchers for automatically detected candidate ripples (events) from 18 channels x 100 events, sorted by agreement rate. Note the large inter-rater variability. **b** Example LFP waveforms of candidate ripples, classified by all raters as ripples (top row, 100% accepted), by 50% (middle row, 50% accepted), and classified by all raters as false-positives (bottom row, 100% rejected). **c** Agreement rate across evaluators for events rated by >50 % of raters as ripples (blue) vs. false-positives (grey). **d** Agreement rate across evaluators for events detected on ripple-negative channels (grey) vs. events detected on ripple-positive channels (blue).

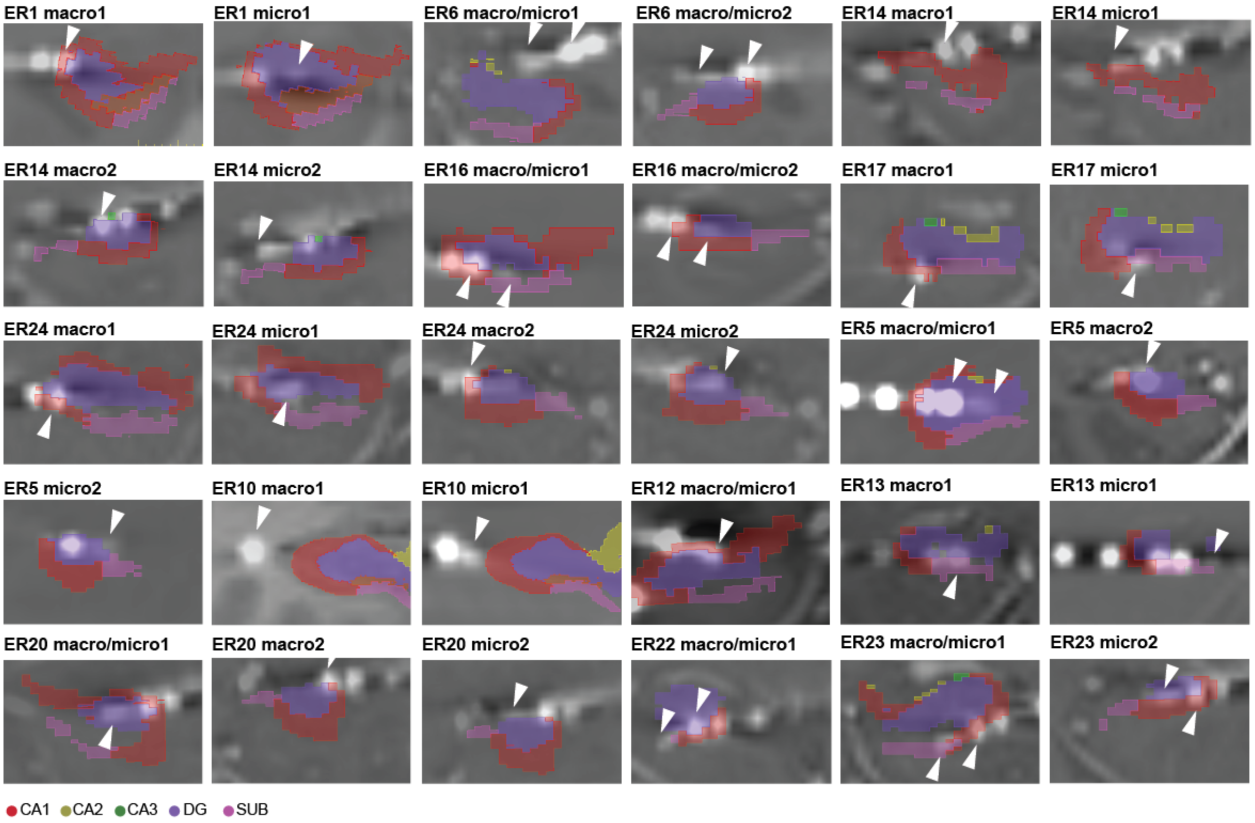


**Supplementary Fig. 4:** Location of human macro/microwire in the hippocampus. Each plot displays a coronal view of the hippocampus, overlapping the patient's postoperative CT-scan with a preoperative T1 scan. The hippocampal subfield segmentation with ASHs is superimposed on the anatomical images to identify the positions of macrocontacts and microwires (white arrowheads). Patient and contact index are identified on top of each plot. Only contacts in the hippocampus are shown.

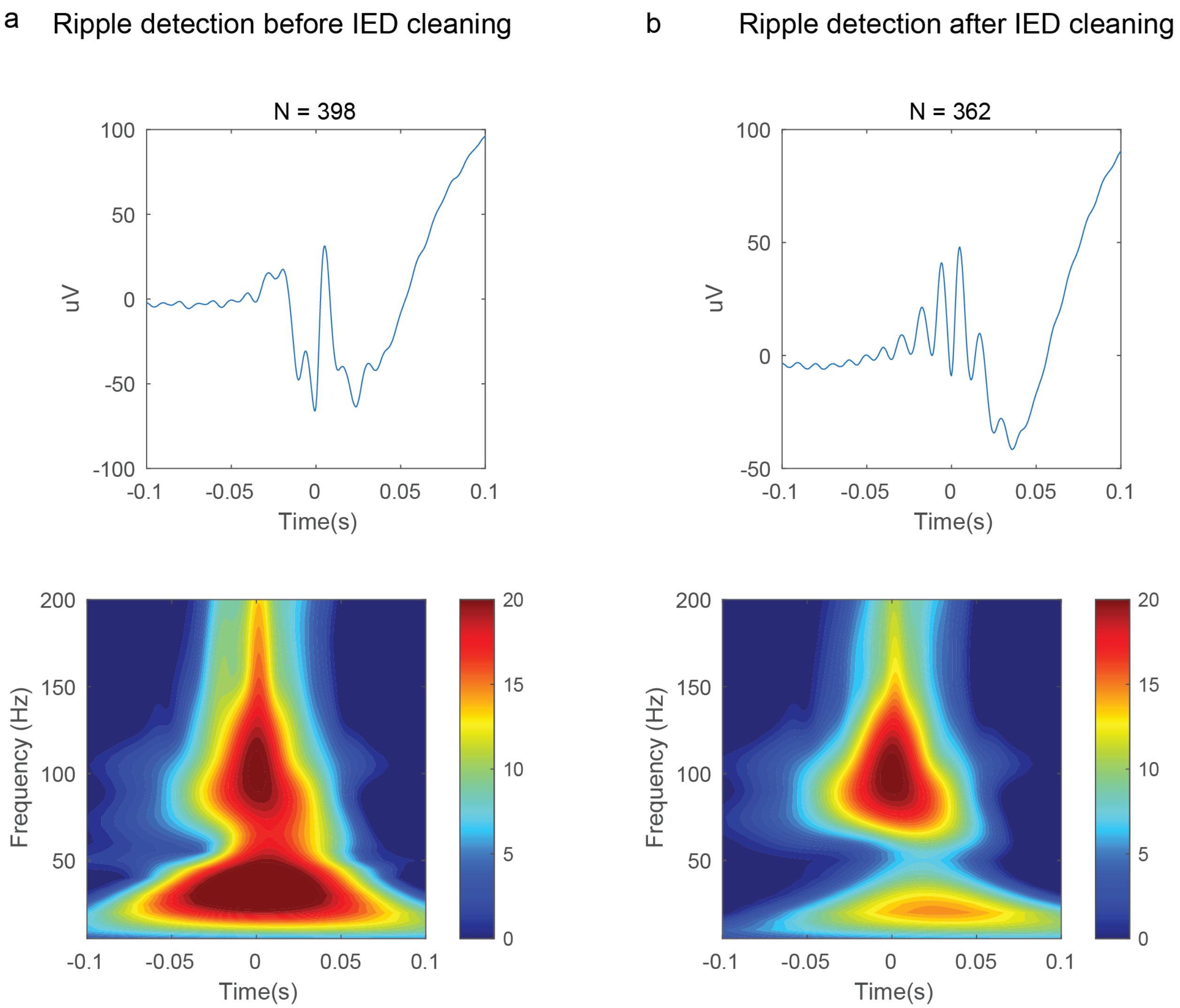


**Supplementary Fig. 5: Discarding IEDs from the raw signal improves the accuracy of ripple detection**. **a** Human average LFP(top) and spectrogram(bottom) of automatically detected ripples from one example human channel. Most of these events were IEDs, obscuring true SPW-Rs. **b** Same as (a) but with removal of periods around IEDs (±1s) from the signal before automatic ripple detection.

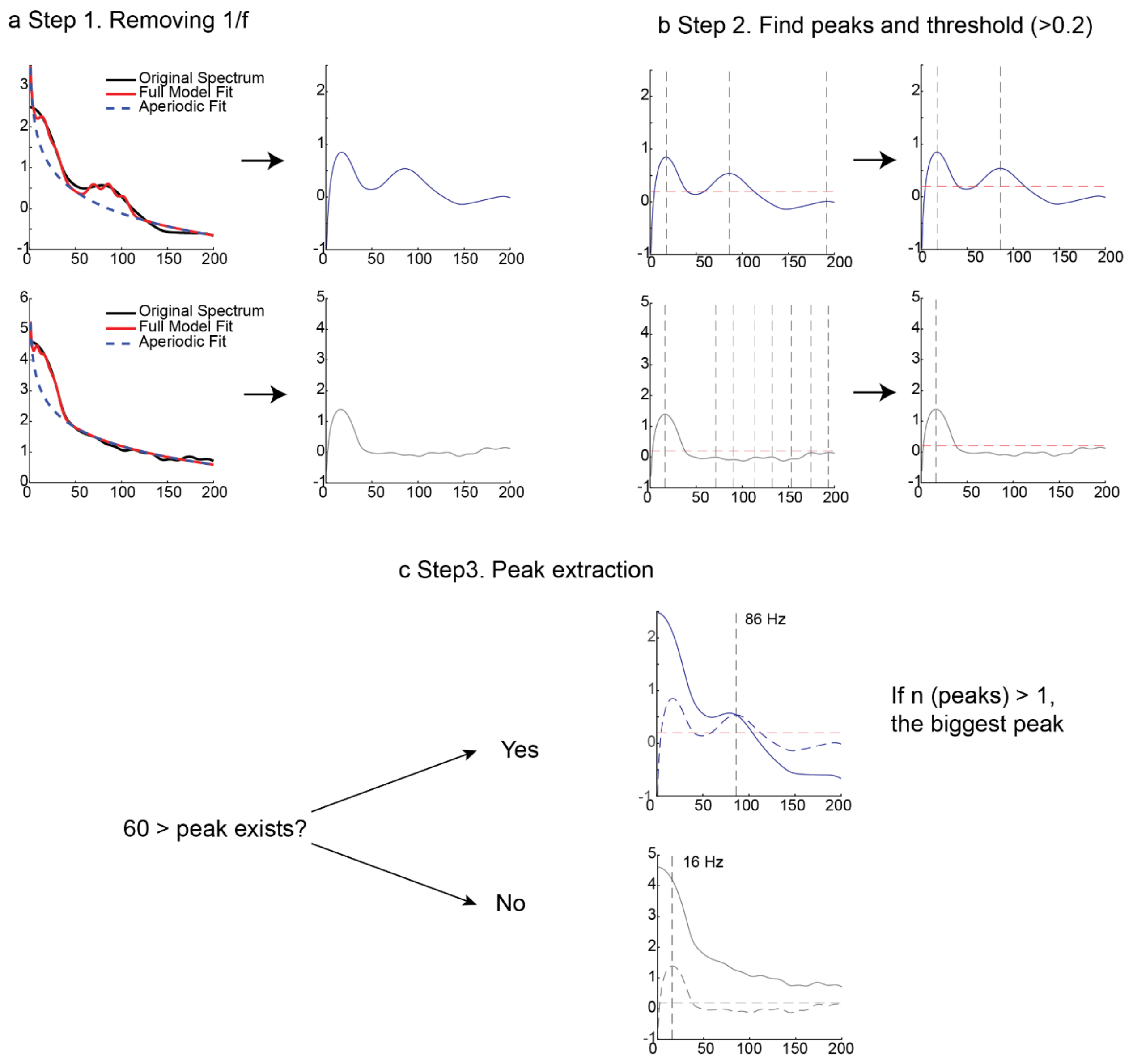


**Supplementary Fig. 6. Spectral peak extraction from 1/f-corrected PSDs. a** A 1/f-corrected PSD was obtained by subtracting the aperiodic component, fitted using FOOOF, from the original spectrum. **b** Peaks were identified from the 1/f-corrected PSD, and only those exceeding the threshold (0.2) were kept. **c** Among the detected peaks, a peak above 60 Hz was selected. If multiple peaks above 60 Hz were present, the largest peak was chosen. When no peak above 60 Hz was identified, the peak at the lowest frequency was selected instead.

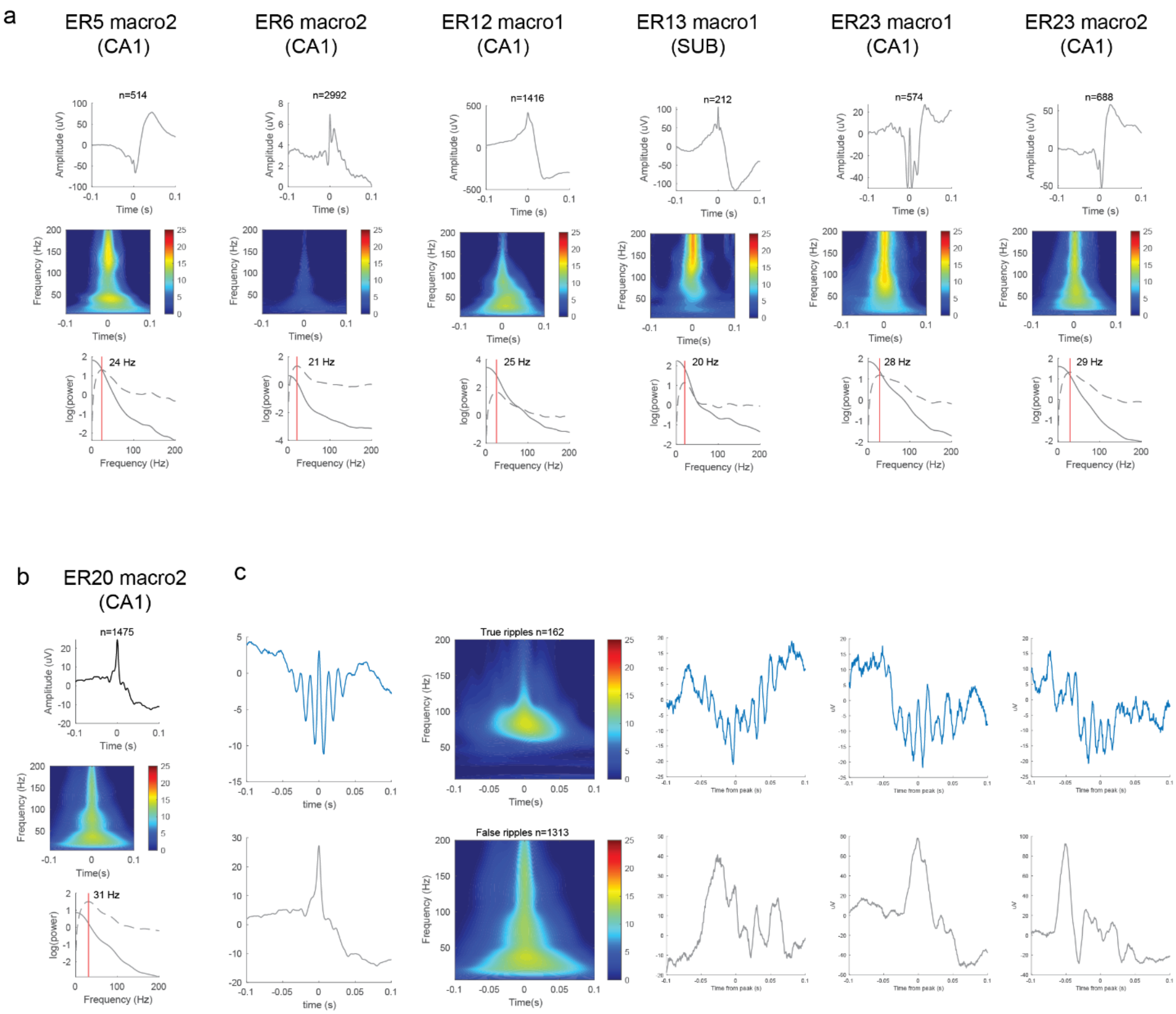


**Supplementary Fig. 7. Human macrocontacts with <60 Hz spectral peak. a** Average LFP waveforms, spectrograms and PSDs of automatically detected candidate ripples on macrocontacts with a spectral peak below 60 Hz. **b** Same as (a) but for the only one ripple positive macrocontact. **c** True ripples (top) and false-positives (bottom) of the same channel as in (b). The true positive rate was 11%. Each row from left to right displays: average LFP waveform of all events, average spectrogram, three example events.

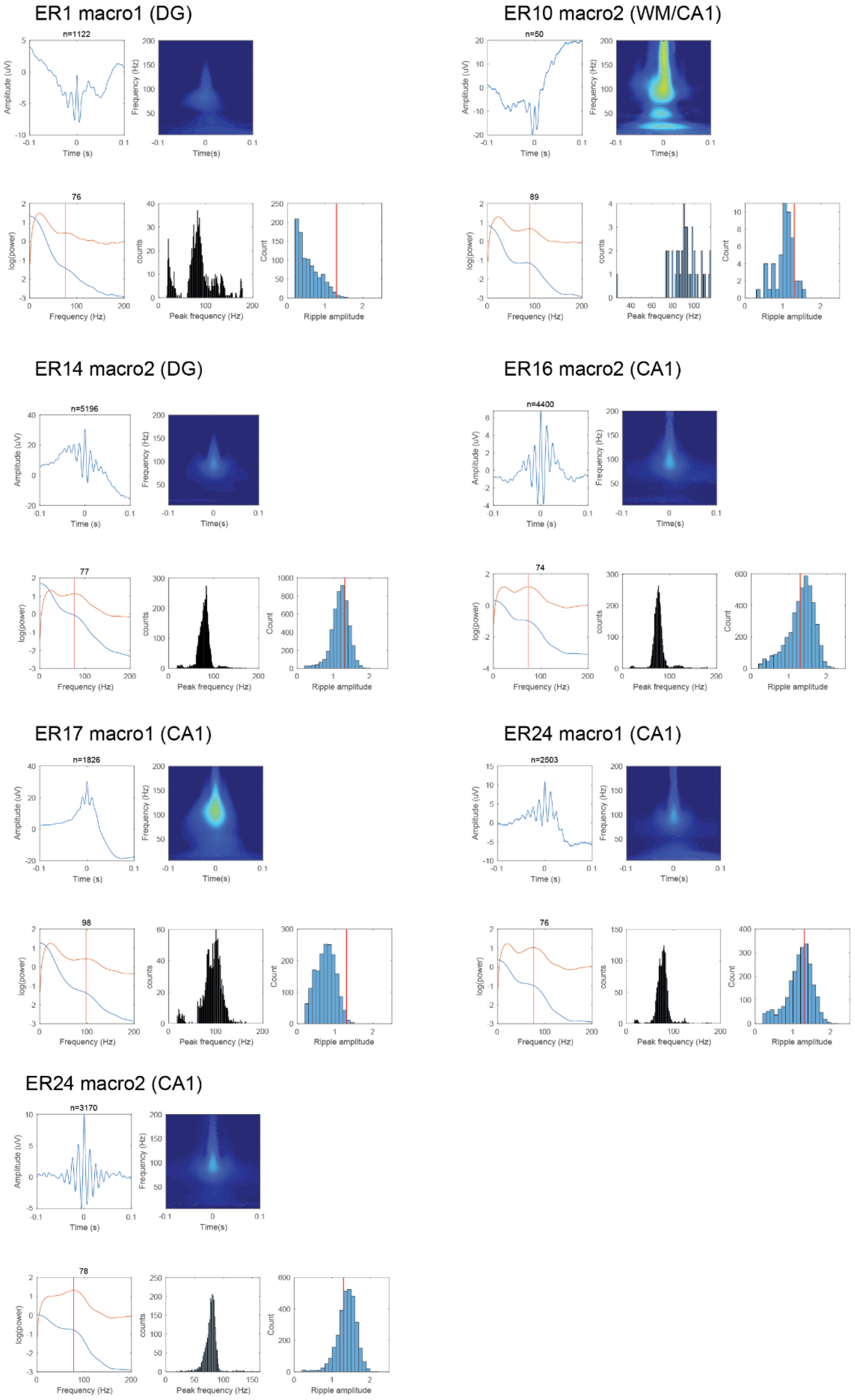


**Supplementary Fig. 8. Human macrocontacts with >60 Hz spectral peak and no ripples found by manual curation**. Average LPF waveform, spectrogram, peri-event PSD, ripple peak distribution and ripple amplitude distribution for each channel. Red lines in the ripple amplitude histograms: mean peak amplitude of true ripples on macrocontacts.

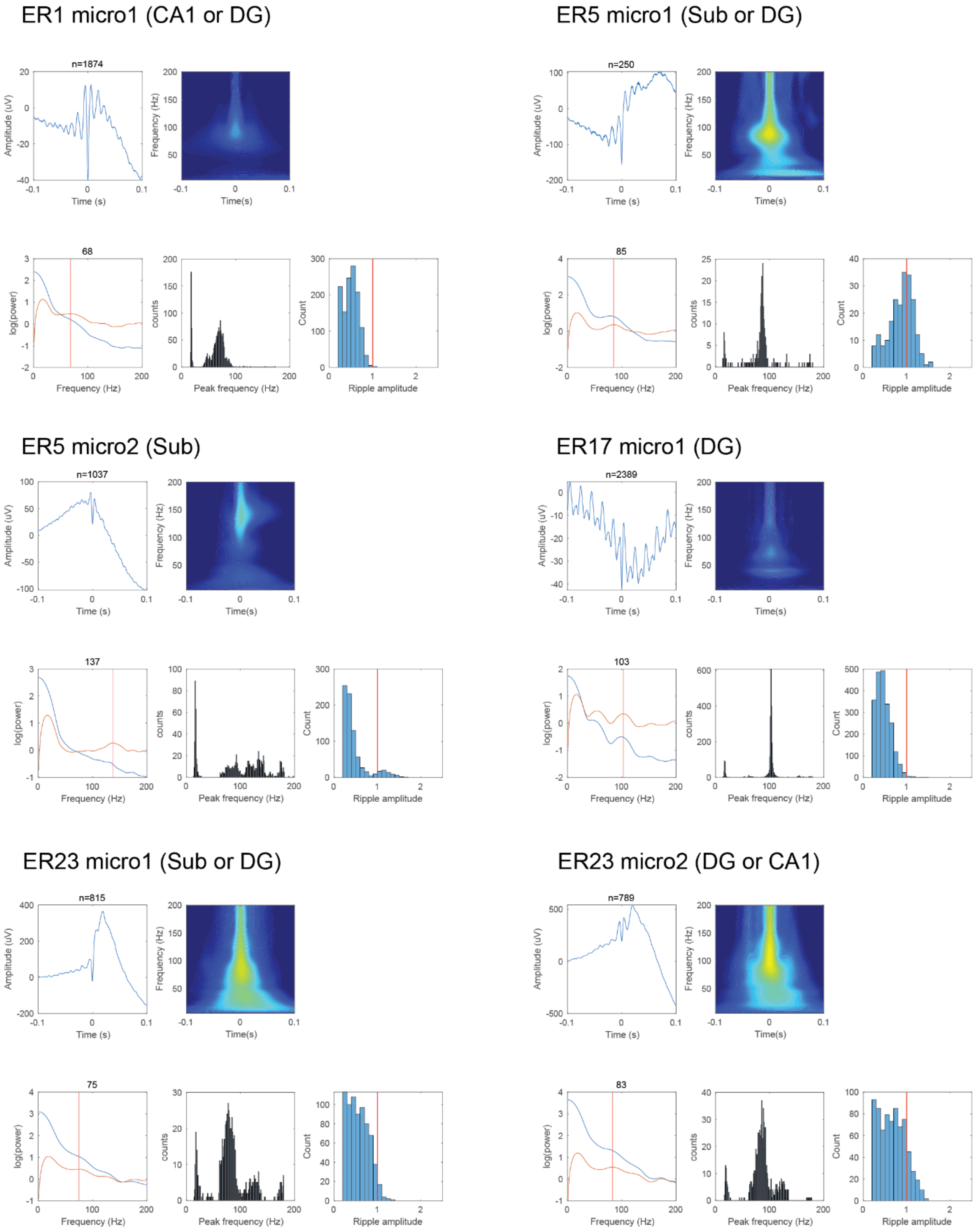


**Supplementary Fig. 9. Human microwire channels with >60 Hz spectral peak and no ripples found by manual curation**. Average LPF waveform, spectrogram, peri-event PSD, ripple peak distribution and ripple amplitude distribution for each channel. Red lines in the ripple amplitude histograms: mean peak amplitude of true ripples on macrocontacts.

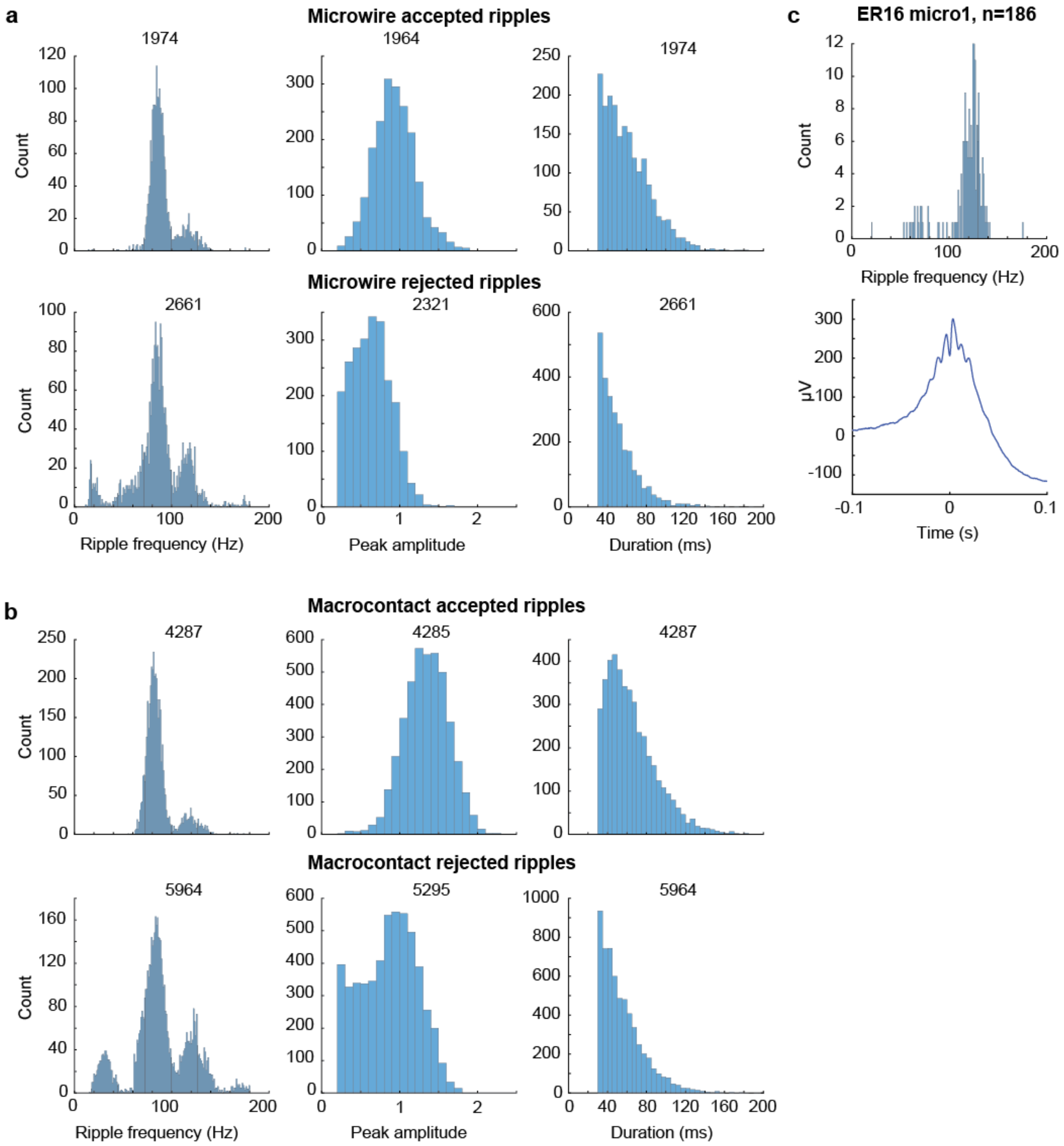


**Supplementary Fig. 10. Human curated ripples are larger and longer than false-positives.** **a** Comparison of human curated ripples (top, accepted) and false-positives (bottom, rejected) on microwires. From left to right: histograms of ripple frequency, peak amplitude, and duration. **b** Same for macrocontacts. **c** Ripple frequency and average LFP of ripples detected on one microwire located in CA1. Positive waveforms suggest location in the pyramidal layer. Note the > 100 Hz ripple frequency distribution.

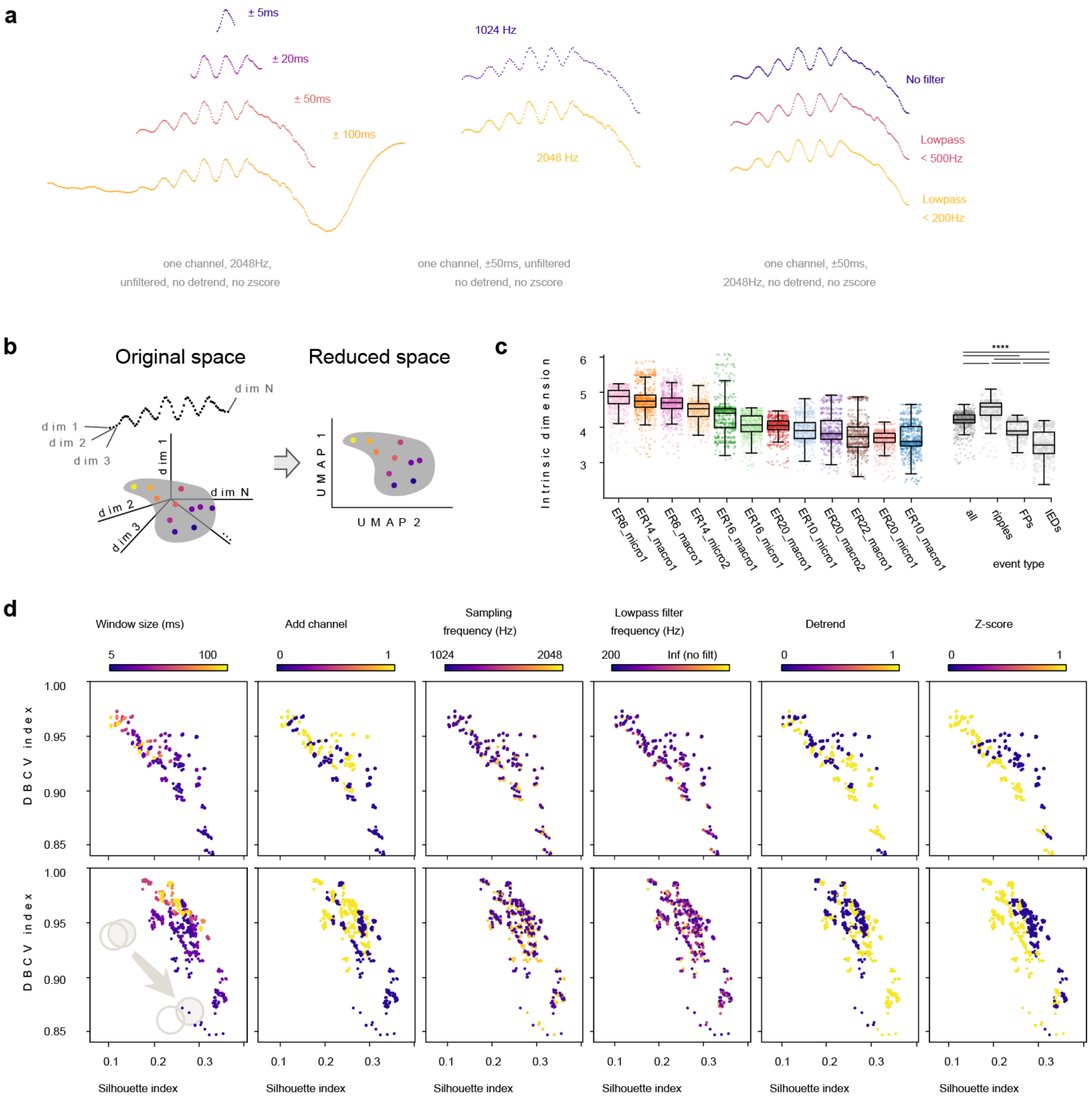


**Supplementary Fig. 11. Topological characterization of events in the original and reduced spaces. a** Examples of how events look like depending on different pre-processing characteristics. **b** Schematic of UMAP dimensionality reduction **c** Computing intrinsic dimension for all possible combinations of parameters (window_size: 5, 10, 15, 20, 25, 50, 75, 100; add_channel: True, False; sampling_frequency: 2048, 1024; lowpass_freq: no filtering, 500, 400, 300, 200; detrend: True, False; zscore: True, False). Checking variability within and between sessions. Comparison of the intrinsic dimension between events labeled as “ripple”, “IED” or “FP” (false positives, or rejected ripples) have significant differences in their dimensionality. FPs lie between the lower dimensionality of IED and higher of ripples. Mean intrinsic dimension of all events is 4D. **d** Mean clustering scores across sessions. We have used DBCV index and Silhouette indexes, very standard indexes to assess similarity between clusters. Low DBCV index and high Silhouette indicates good clustering. Here, each point represents the mean DBCV and Silhouette indices between the IED and ripple events of all sessions, for a particular set of pre-processing parameter combination. Upper row shows the metrics computed in the original space, lower row in the reduced space. Reducing the dimensionality preserves the structure of the mean scores.

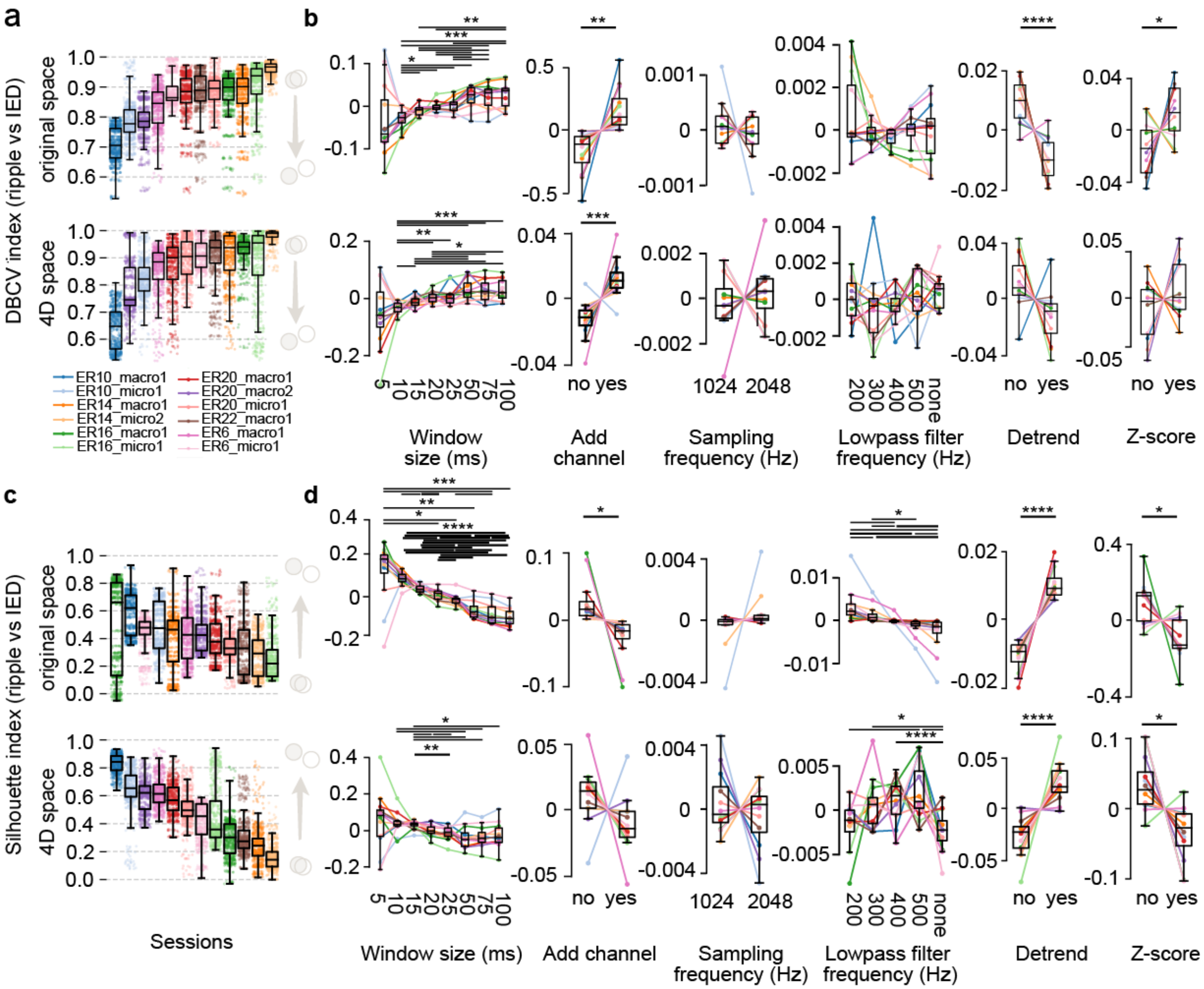


**Supplementary Fig. 12. Snippet variables that optimize ripple-IED segregation**

**a.** DBCV index to evaluate IED-ripple clusterization per session, both in the original space (top) or the 4D space (bottom). Sessions have very different DBCV scores, indicating different levels of curation potential. **b.** DBCV index changes depending on event features, in the original space (top) and the 4D space (bottom). Each line depicts the median of the cluster metric for a single patient for all possible values of the rest of the parameters. Due to the variability in the mean DBCV index across session, the mean score per session has been subtracted. Tendencies remain equal for the 4D space. Best parameters are: small windows (5-10ms), no extra channels, 400-500 Hz lowpass filter, detrending and no z-scoring. Sampling frequency is not significant. **c,d.** Same as a,b but for the Silhouette index. Same conclusions.

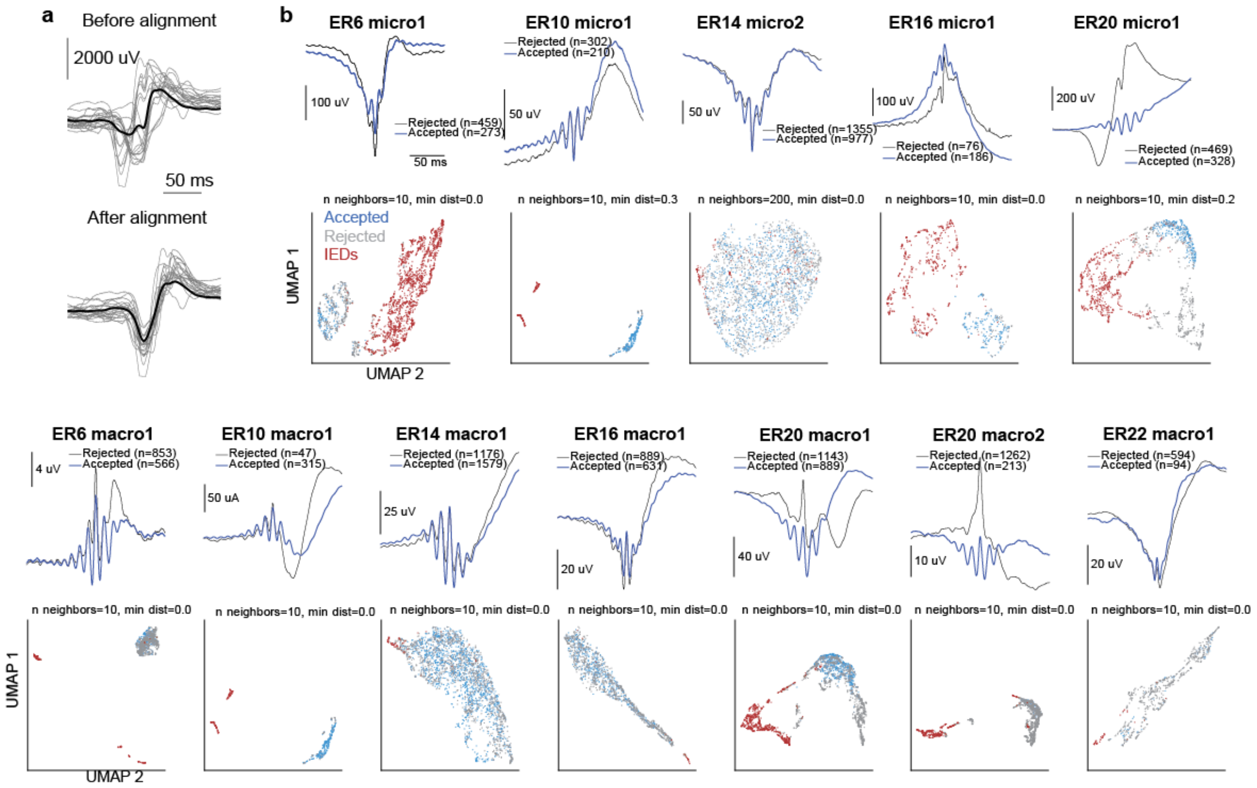


**Supplementary Fig. 13. UMAP embedding of events detected on all human ripple-positive channels. a.** Aligning IED peaks. **b.** UMAP embedding of IEDs and ripples. For each channel top plots display average waveforms of manually curated (accepted) ripples (blue) and manually labelled rejected events (rejected,grey). Bottom plots: Common UMAP embedding of all candidate ripples (blue – accepted, grey -rejected) and automatically detected IEDs (red).

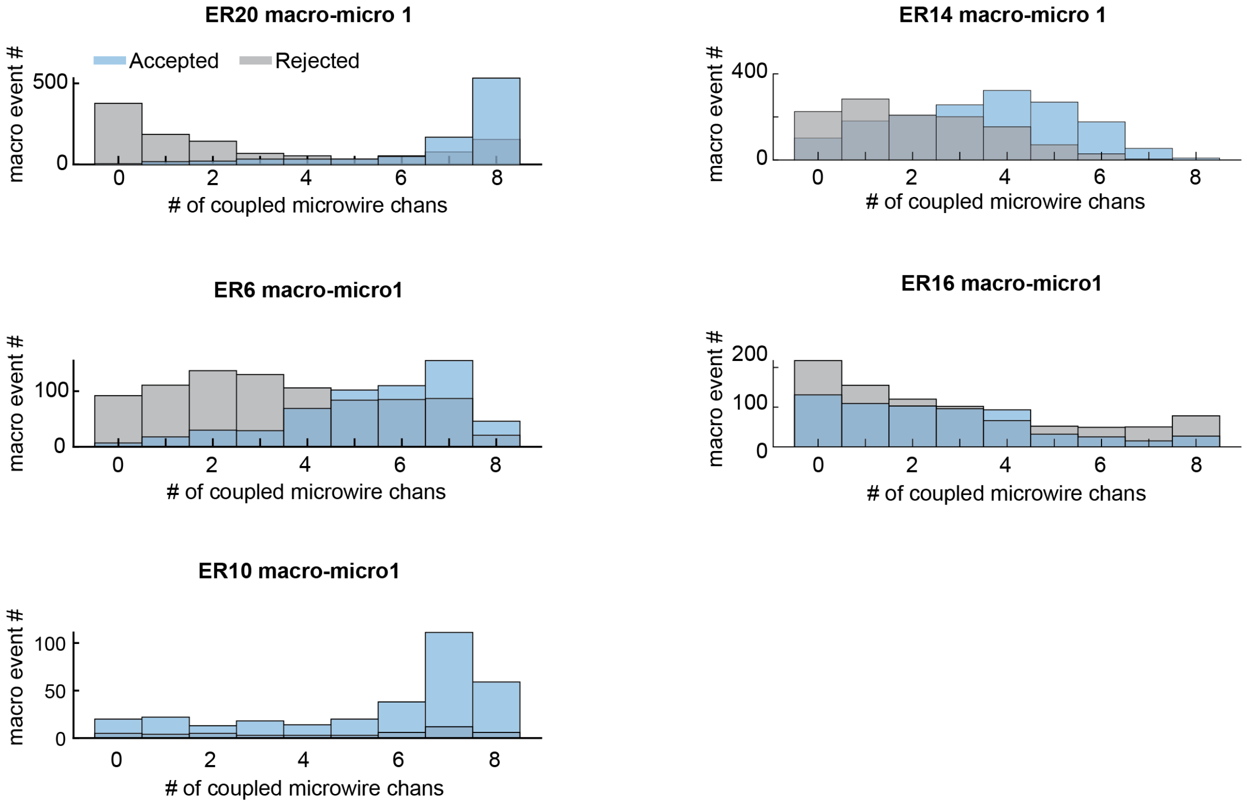


**Supplementary Fig. 14**. **Number of coupled microwire channels during accepted and rejected macrocontact events.** Blue: Accepted events, Gray: Rejected events. Note that rejected events are less correlated with microwire ripple activities.

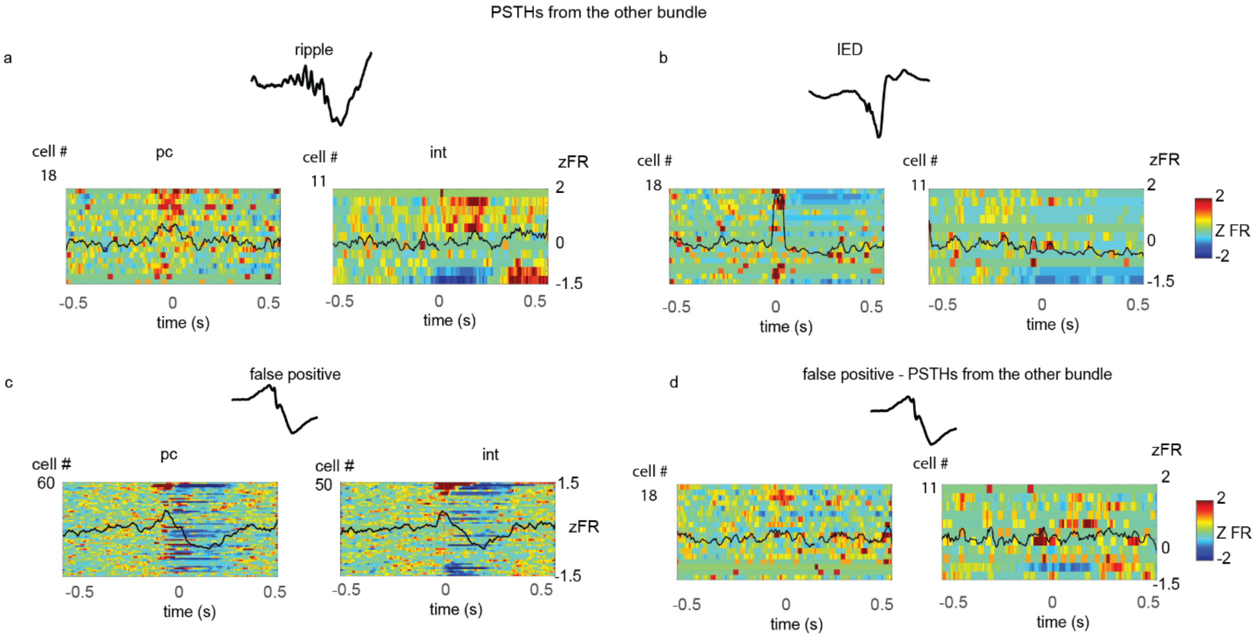


**Supplementary Fig. 15 Peri-event PSTHs of human ripples and IEDs on remote microwire bundles and of false-positive events. a** Z-scored peri-ripple spiking in principal cells (pc) and interneurons (int) in patients implanted with >1 microwire bundles. Only neurons recorded from microwires on the remote depth electrode from the one used for ripple detection are shown. Neurons are sorted by strength of peri-ripple positive neuronal modulation. Black traces: mean z- scored firing rates of all cells around ripples. Color axis: z- scored firing rates [−1.5 to 2]. **b** same as (a) for IEDs. **c** peri-event spiking rates for false-positives (defined by manual curation) of automatically detected ripples. Neurons recorded from microwires within the same depth electrode used for ripples detection are shown. Black traces: mean z- scored firing rates. **d** same as (c) for neurons detected on microwires within the remote depth electrode from the one used for ripples detection.
